# Loss of TCA cycle turning promotes stem cell function

**DOI:** 10.64898/2026.09.21.753355

**Authors:** Yafeng Li, Joseph Rose, Edward O. Kwarteng, Xiaotong Yang, Jacob Zielke, Li Li, Zhiyu Zhao, Ji Hyung Jun, Michalis Agathocleous

## Abstract

The citric acid cycle (TCA cycle) is the common terminal pathway for the oxidation of all nutrients. Citrate oxidation to oxaloacetate produces CO_2_, and citrate synthase (CS) uses nutrient-derived acetyl groups to regenerate citrate and fuel cycle turning. However, the essentiality of cycle fueling and turning *in vivo* remains unclear. Here, we use hematopoiesis, the most proliferative system in the body, as a model to show that, contrary to common assumptions, TCA cycle turning is dispensable for respiration, survival, and proliferation of stem and progenitor cells *in vivo* and its loss promotes stem cell function. Hematopoietic-specific *Cs* deletion in adult mice blocked citrate cycling without reducing the frequency of hematopoietic stem (HSC) and progenitor cells. HSCs and progenitor cells adapted to TCA cycle loss by markedly increasing nutrient consumption and biosynthesis. Disruption of cycle turning increased HSC regeneration, myeloid progenitor proliferation, and myelopoiesis *in vivo.* HSCs without a turning TCA cycle outcompeted wild-type HSCs within the same environment. The effect of CS deletion on HSC function was not phenocopied by genetic ablation of cytosolic citrate use and was rescued by ablation of glutamine use in biosynthesis. Therefore, TCA cycle turning restrains nutrient uptake, biosynthesis, cell proliferation, and stem cell function. These results suggest an explanation for the reduction in cycle activity observed in many normal proliferating cells and cancer cells.

## Introduction

The complete oxidation of all nutrients to CO_2_ is accomplished by a single terminal pathway, the citric acid cycle (or tricarboxylic acid, TCA cycle). The cycle burns two carbons per turn by converting 6-carbon citrate to 4-carbon oxaloacetate and regenerates citrate from nutrient-derived 2-carbon acetyl groups in the citrate synthase (CS) reaction^1^. CS delivers the fuel that keeps the cycle turning^1,2^ (**Fig. 1a**). The discovery of citrate regeneration from oxaloacetate and a triose-derived molecule in the CS reaction led Krebs to formulate the concept of the cycle as the central catabolic engine^1,3,4^. Nearly a century of work has enshrined this pathway in textbooks as an integral part of oxidative energy production. However, it has been challenging to determine if respiration or other aspects of cellular metabolism require the closed-circuit, turning TCA cycle *in vivo,* or indeed if most cells require the cycle’s turning for survival and proliferation, because the reactions from citrate to oxaloacetate also participate in linear offshoots from the cycle, whose disruption can block biosynthesis^5,6^. In principle, CS deletion can selectively block cycle turning but preserve the pathway between citrate and oxaloacetate and thus dissociate the cycling pathway from its linear offshoots **(Fig. 1a)**. But CS germline deletion is embryonically lethal, and conditional deletion in specific cell types has not been tested. Thus, the extent to which cells *in vivo* require a fully turning TCA cycle to survive remains unclear.

**Figure 1.**
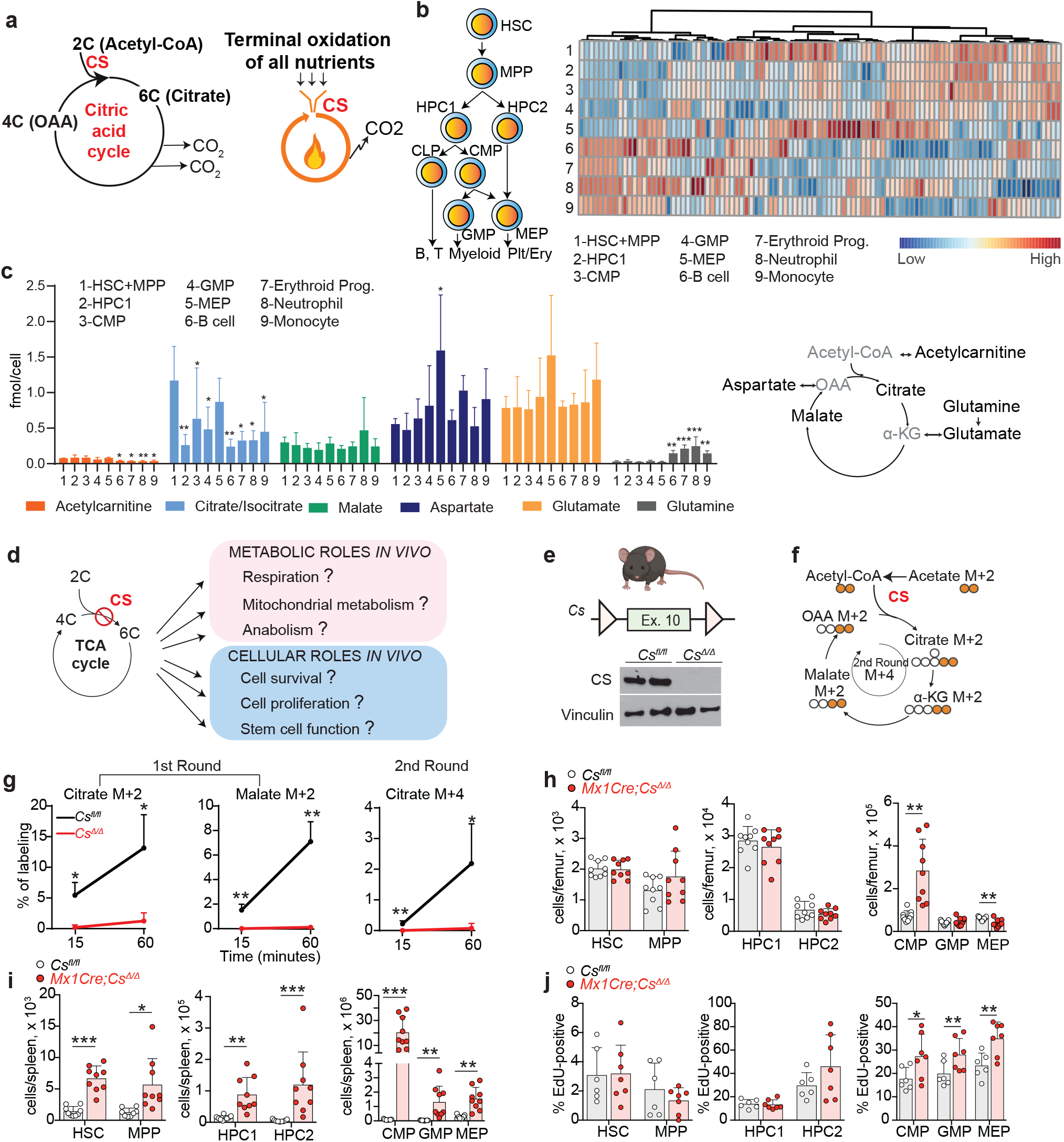
The TCA cycle is dispensable for HSPC survival and proliferation *in vivo*. a. CS funnels acetyl groups derived from the catabolism of all nutrients to the TCA cycle for terminal oxidation. Left: The TCA cycle converts citrate to oxaloacetate and regenerates citrate from oxaloacetate *via* CS. Right: CS is the key to the terminal oxidation of all nutrients. b. Metabolomics analysis of hematopoietic differentiation showing that each cell population has a distinct metabolite composition (abbreviations for each cell population and markers used for isolation are described in the methods. Additional differentiation routes may operate along with the ones shown in the differentiation tree). c. Absolute levels of TCA cycle-related metabolites in the hematopoietic cell types noted in (b). * indicates comparison with HSC+MPP (n = 5 mice). d. Possible metabolic and cellular consequences of disrupting CS. e. Gene targeting design for *Cs^fl^* mice and western blot showing CS depletion in *Mx1Cre;Cs^Δ/Δ^* bone marrow cells. f. Illustration of U^13^C-acetate tracing to assess the first 1^st^ and 2^nd^ round of TCA cycle turning *ex vivo*. g. Labeling of TCA cycle metabolites from U^13^C-acetate in bone marrow cells of *Mx1Cre;Cs^Δ/Δ^* mice and *Cs^fl/fl^* littermate controls (n = 4-5 mice/genotype). h-i. Number of HSCs and progenitors in the bone marrow and spleen of *Mx1Cre;Cs^Δ/Δ^* mice and *Cs^fl/fl^* littermate controls 3 weeks after deletion. j. Proportion of EdU-positive cells after a 2-hour EdU pulse within different bone marrow HSPC populations 3 weeks after deletion. All data represent mean ± s.d. Statistical significance was assessed with a one-way ANOVA of log-transformed data (c), or a Mann-Whitney test (g-citrate M+2 15’, malate M+2 15’, citrate M+4, i-HPC2, CMP), or a Welch’s test (g-citrate M+2 60’, malate M+2 60’, h, i), or a t-test (j). All figures show * 0.01<p<0.05, ** 0.001<p<0.01, ***p<0.001. Statistical tests were chosen after testing for assumptions of normality and equality of variance as detailed in the Methods section. Individual datapoints in all dot plot graphs in the paper represent biological replicates (typically, mice).

It is also unclear which aspects of cellular metabolism require a turning TCA cycle *in vivo*. Classically, the cycle supports aerobic respiration by linking nutrient oxidation to the electron transport chain to convert nutrient energy to ATP^3,6^. It may also supply anabolic intermediates for cell proliferation^3,6,7^. However, nutrient oxidation is reduced in many proliferating stem or progenitor cells as compared to differentiated cells^8–10^. Similarly, most examined cancers have slower TCA cycle turning as compared to normal tissues *in vivo*^11,12^. These findings raise the question of the role of TCA cycle turning in proliferating cells *in vivo*.

The highest proliferative activity in the body is found in the bone marrow, where hematopoietic cells produce more cells than all other tissues^13,14^. Hematopoietic cells are derived from self-renewing hematopoietic stem cells (HSCs). HSCs have low levels of oxidative metabolism compared to progenitor cells^10,15–22^. Paradoxically, deletion of TCA cycle components succinate dehydrogenase^23^ or fumarate hydratase (FH)^24^ or of complex III of the electron transport chain (ETC)^25^ impairs HSC self-renewal or differentiation, suggesting HSCs require aspects of oxidative metabolism. In addition to impacting the TCA cycle, these manipulations also block biosynthetic reactions from glutamine to aspartate, inhibit the electron transport chain^23,25^, or elevate succinate, fumarate, or 2-hydroxyglutarate, which exert epigenetic effects that impact HSC function^24,25^. It thus remains unclear if TCA cycle turning is essential for *in vivo* survival and proliferation of HSCs or other somatic stem cells. The cyclic pathway forms the terminal step for the complete oxidation (combustion) of all nutrients. *Cs* deletion provides a way to directly interrogate the role of cycle turning in proliferating cells *in vivo*.

## RESULTS

### Metabolic differentiation in hematopoiesis

Metabolism has been mostly analyzed in cultured cells or whole tissues, limiting our understanding of metabolic heterogeneity among cell types *in vivo*. The marrow offers unique advantages to approach this problem because highly purified cell types can be isolated at low temperature without enzymatic dissociation. To study cell-type-specific metabolism *in vivo*, we used rare cell metabolomics methods we developed^26,27^ to construct an expanded^26^ metabolomic map of hematopoiesis (**Fig. 1b**). We pooled CD150^+^CD48^-^Lin^-^Sca-1^+^Kit^+^ HSCs and CD150^-^CD48^-^Lin^-^Sca-1^+^Kit^+^ MPPs because we previously showed that these cell types are almost identical in their metabolome^26^. Each hematopoietic cell type was marked by unique metabolic features (**Fig. 1b, Supplementary Table 1**) consistent with our previous work^26,27^. To specifically measure absolute levels of TCA cycle-related metabolites, we used a spiked-in mixture of stable isotope-labeled internal standards (**Fig. 1c**). Relative to differentiated cells, HSCs and progenitors (HSPCs) were characterized by low glutamine and high citrate and acetylcarnitine, which reflects acetyl-CoA levels. Analysis of publicly available gene expression data showed that TCA cycle-related enzymes had cell-type-specific expression patterns in hematopoietic differentiation, with overall decline in expression at late stages of myeloid and lymphoid differentiation and an increase in expression in erythroid lineage progenitors (**Extended Data Fig. 1a-b)**. These data suggest remodeling of the TCA cycle with hematopoietic differentiation.

### TCA cycle turning is dispensable for HSC and progenitor survival and proliferation

To test if TCA cycling is required for HSPC metabolism, survival, and function *in vivo* (**Fig. 1d**), we developed a *Cs* conditional knockout mouse. *Cs^fl^* mice were generated by floxing exon 10 to ablate CS amino acids 340-409, which include part of the active site (**Fig. 1e**). *Cs* was conditionally deleted in adult hematopoiesis using *Mx1Cre.* CS was depleted in the bone marrow 3 weeks after poly I:C-mediated deletion (**Fig. 1e**). To validate that *Cs* deletion prevents cycle turning, *Cs^Δ/Δ^* or littermate control hematopoietic cells were incubated *ex vivo* with U^13^C-acetate, which fuels the cycle with acetyl groups (**Fig. 1f**). *Cs* deletion abolished U^13^C-acetate-derived labeling from the first and second round of the cycle in citrate/isocitrate (**Fig. 1g**) and other TCA cycle-related metabolites (**Extended Data Fig. 1c-d**), indicating a complete loss of cycle turning.

To test if the cycle is required for cell survival *in vivo,* we analyzed *Cs-*deficient hematopoietic cells 3 weeks after deletion. Genotyping of colonies derived from single sorted *Cs^Δ/Δ^*HSCs or CD48^+^Lin^-^Sca-1^+^Kit^+^ oligopotent hematopoietic progenitors (HPCs) confirmed that 98-100% of HSPCs were deleted (**Extended Data Fig. 2a**). CS was depleted in purified HSCs or progenitors (**Extended Data Fig. 2b**). *Cs* deletion did not impact the number (**Fig. 1h**), and increased the frequency of bone marrow HSCs, MPPs, and CD150^-^CD48^+^Lin^-^Sca-1^+^Kit^+^ HPC1 cells (**Extended Data Fig. 2c, d**). *Cs* deletion increased the frequency or number of spleen HSCs, MPPs, and HPCs, and of bone marrow and spleen myeloid progenitors (**Fig. 1h,i, Extended Data Fig. 2e-h**). The TCA cycle may supply anabolic intermediates for proliferation^3,6,7^. However, *Cs* deletion did not impair the proliferation of slowly dividing HSCs, MPPs or of more rapidly dividing HPCs (**Fig. 1j)**. CMPs, GMPs, and MEPs divide 1-2 times/day and are among the most rapidly dividing cells in the body^28,29^. *Cs* deletion increased their proliferation (**Fig. 1j**).

*Cs^Δ/Δ^* mice had a hypocellular bone marrow and splenomegaly (**Extended Data Fig. 2i-j**). *Cs* deletion blocked marrow erythropoiesis at the CD71^+^Ter119^+^ erythroblast stage causing compensatory spleen erythropoiesis and anemia (**Extended Data Fig. 2k-m).** *Cs^Δ/Δ^* deletion preserved platelets and most myeloid cells, except for decreased marrow and increased spleen neutrophils (**Extended Data Fig. 2n-q).** *Cs* deletion reduced bone marrow and blood B cells and impaired B cell development (**Extended Data Fig. 2r-t**). Most effects of *Cs* on hematopoiesis persisted at 5 weeks post-deletion (**Extended Data Fig. 3a-v**). At this time, *Cs^Δ/Δ^* thymic T cell progenitors and splenic T cells were also reduced (**Extended Data Fig. 3w-y**). *Mx1Cre;Cs^Δ/Δ^* mice died 6-7 weeks after deletion with anemia and lymphopenia but normal myeloid and platelet cell numbers (**Extended Data Fig. 3z-e’**). Therefore, CS is required for lymphopoiesis and erythropoiesis, but is dispensable for HSC or progenitor survival, proliferation, myeloid and platelet differentiation (**Extended Data Fig. 3f’**).

### TCA cycle turning is dispensable for aerobic respiration

The classical function of the cycle is to extract energy stored in nutrients by linking nutrient catabolism to respiration^3,6^. Terminal nutrient catabolism in the cycle to CO_2_ produces reducing equivalents used in the ETC to produce ATP. To test if the TCA cycle is needed for respiration, we assessed mitochondrial metabolism in *Cs-*deficient cells. *Cs* deletion did not reduce respiration in total bone marrow hematopoietic cells and, surprisingly, increased respiration in freshly sorted HSPCs (**Fig. 2a, Extended data Fig. 4a-b**). Therefore, TCA cycle turning is dispensable for respiration in hematopoietic cells. ATP levels declined in *Cs-*deficient HSC+MPP and HPC1 and did not change in myeloid progenitors or total hematopoietic cells (**Fig. 2b**). NADH levels declined in *Cs-*deficient HSC+MPP and NAD^+^ levels did not change (**Fig. 2c)**. Because respiratory O_2_ consumption increased in *Cs^Δ/Δ^* HSPCs, reduced ATP and NADH levels may reflect increased utilization or decreased production. *Cs* deletion decreased the mitochondrial membrane potential in some HSC and progenitor cell types but not others and did not reduce mitochondrial mass (**Extended data Fig. 4c-d**). Oxidative metabolism produces reactive oxygen species (ROS); however, *Cs* deletion did not change ROS levels in HSCs or progenitor cells (**Extended data Fig. 4e**). These results suggest that the TCA cycle helps to maintain the mitochondrial membrane potential and NADH levels of HSCs and progenitors, but it is not needed to fuel respiration.

**Figure 2.**
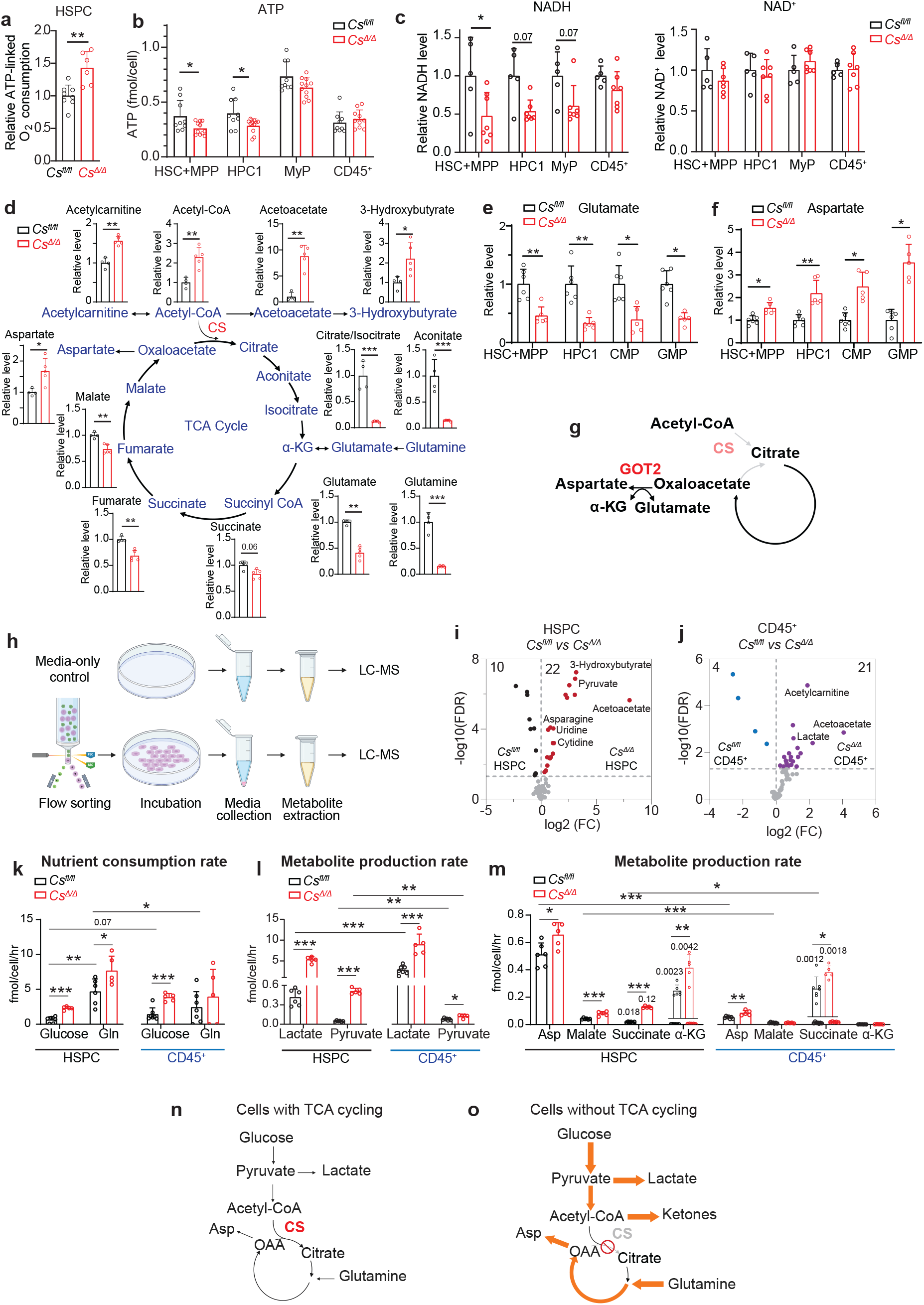
The TCA cycle is dispensable for respiration and its loss reprograms the metabolome and increases total metabolite production and nutrient consumption. a-c. Respiratory oxygen consumption (a), ATP levels (b), or NADH and NAD^+^ levels (c) of sorted Lin^-^kit^+^ HSPCs or CD45^+^ total bone marrow cells from *Mx1Cre;Cs^Δ/Δ^* mice or *Cs^fl/fl^* littermate controls. d. Metabolomics analysis of total BM cells. e-f. Glutamate and aspartate levels of sorted HSPC populations. g. CS and GOT2 compete for oxaloacetate. h. Design of the nutrient consumption and metabolite production experiment. i-j. *Cs^Δ/Δ^* cells overproduce a greater number of metabolites than wild-type cells. k. Absolute quantification of the rate of glucose and glutamine consumption. l-m. Absolute quantification of the rate of metabolite production. n-o. Metabolic rewiring after *Cs* deletion. All data represent mean ± s.d. Statistical significance was assessed with a t-test (a, b-HSC+MPP), Mann-Whitney test (b-HPC1), or multiple t-tests of log-transformed metabolomics data with multiple comparisons correction by controlling the false discovery rate at 5% with the Benjamini, Krieger, and Yekutieli method (d-f), or in k-m with Welch’s t-test (HSPC-malate, pyruvate, lactate, glucose vs. glutamine, HSPC vs. CD45^+^-aspartate), Mann-Whitney test (CD45^+^-pyruvate, HSPC vs CD45^+^-glucose, pyruvate), or t-test (other comparisons).

### Loss of TCA cycle turning causes widespread metabolic reprogramming

To comprehensively understand the metabolic roles of TCA cycling *in vivo* we analyzed the metabolome of bone marrow cells from *Mx1Cre;Cs^Δ/Δ^* or littermate control mice. Of 323 detected metabolites, the levels of 71 metabolites increased, and the levels of 71 decreased (**Extended Data Fig. 4f-h, Supplementary Table 2**), suggesting that cells without a TCA cycle reprogram a large part of their metabolism. *Cs* deletion reduced the levels of some TCA cycle-related metabolites, including citrate/isocitrate, aconitate, glutamate and glutamine and increased the levels of upstream metabolites acetylcarnitine, acetyl-CoA, and ketone bodies (**Fig. 2d**). This suggested redirection of acetyl groups not burnt in the cycle to ketones^2,30^. *Cs* deletion elevated metabolites in lower glycolysis or its branching biosynthetic pathways (**Extended Data Fig. 5a**) and impacted several other pathways, including amino acid and nucleotide metabolism (**Extended Data Fig. 4h, 5b**). *Cs* deletion increased the levels of most amino acids and of many dipeptides, except for glutamate and glutamine (**Extended Data Fig. 5c-d**). The increased levels of carbohydrate and protein catabolic intermediates in *Cs*-deficient cells support the idea that a major metabolic role of the TCA cycle is to remove excess nutrient-derived metabolites^2,31^.

A common but rarely addressed problem in metabolomics is that metabolic changes at the tissue level may reflect changes in the composition of metabolically dissimilar cell types, rather than changes in the metabolism within individual cell types. To circumvent this, we isolated the three most abundant bone marrow cell types, neutrophils, B cells, and erythroid progenitors and profiled their metabolome. Many of the metabolic changes observed in *Cs-*deficient bone marrow were also observed in these cell types, including elevated acetylcarnitine and aspartate, and depleted glutamine and glutamate (**Extended Data Fig. 6a-c**). These results define core metabolic roles of the TCA cycle *in vivo* that are common across cell types.

To test how HSC metabolism is affected by *Cs* deletion, we used metabolomics for rare cells^26,27^. We and others showed that the levels of most detected metabolites remain stable during hematopoietic cell purification^26,27,32^. *Cs* deletion changed the levels of several metabolites in HSCs and progenitors (**Extended Data Fig. 6d**, **Supplementary Tables 3-6**), suggesting that TCA cycle blockade rewires HSC and progenitor metabolism. *Cs-*deficient HSCs+MPPs or progenitors accumulated aspartate and depleted glutamate (**Fig. 2e-f**). Aspartate and glutamate are connected to CS via the glutamate/oxaloacetate (GOT2) reaction, which competes with CS for oxaloacetate, consumes glutamate, and produces aspartate (**Fig. 2g**). In the absence of CS, oxaloacetate is available for aspartate synthesis. Aspartate synthesis by GOT2 promotes protein and nucleotide synthesis in HSCs and other cells^33,34^, suggesting that loss of TCA cycle turning allows 4-carbon units to be redirected to biosynthesis.

### Loss of TCA cycle turning increases nutrient consumption

To understand how *Cs-*deficient HSPCs maintain or increase their respiration rate (**Fig. 2a**) when they cannot completely oxidize nutrients, we assessed the rates of net metabolite production and consumption by freshly sorted HSPCs or total CD45^+^ bone marrow cells (**Fig. 2h**). HSPCs had distinctive metabolite production patterns compared to total bone marrow (**Extended Fig. 7a-b, Supplementary Table 7**). Of a total 74 produced metabolites, 46 were produced at higher levels by HSPCs, and only 10 by total bone marrow (**Extended Fig. 7c**), suggesting HSPCs have increased metabolic diversity compared to their differentiated counterparts. *Cs* deletion changed the metabolic production landscape (**Extended Fig. 7a-b)** and caused 2-fold more (22 *vs* 10) metabolites to be overproduced than underproduced in HSPCs, and 5-fold more (21 *vs* 4) to be overproduced in total bone marrow (**Fig. 2i-j, Extended Fig. 7d-e**). Some of these metabolites were common in HSPCs and total bone marrow but about half were specific to HSPCs (**Extended Data Fig. 7d-j**). These results provide the first systematic analysis of metabolic end-products of hematopoietic stem cells. They suggest that CS loss increases overall metabolite production.

We then measured the absolute rates of glucose and glutamine consumption and production of major metabolites. Despite the prevailing idea that HSPCs are glycolytic^10^, they consumed a similar or lower amount of glucose, produced less lactate (**Fig. 2k-l**) and consumed more glutamine than total bone marrow cells (**Fig. 2k**). *Cs* deletion increased glucose consumption. This quantitatively corresponded to increased lactate production, consistent with elevated glycolysis (**Fig. 2k-l**). *Cs* deletion further increased HSPC glutamine consumption (**Fig. 2k**). Glutamine is partially oxidized in the linear pathway downstream of α-ketoglutarate to yield 1 CO_2_ and 4-carbon metabolites. *Cs* deletion increased HSPC net production of α-ketoglutarate, malate, succinate, and aspartate, suggesting partial glutamine oxidation (**Fig. 2m**). *Cs^Δ/Δ^* HSPCs overproduced ketone bodies (**Extended Data Fig. 7a,d,f**), suggesting that in the absence of TCA cycle turning, carbon is diverted to ketones. *Cs*-deficient HSPCs overproduced amino acids, including alanine and asparagine, and nucleosides consistent with increased biosynthetic pathway activity (**Extended Data Fig. 7a,d,g,h**). Therefore, *Cs*-deficiency leads HSPCs to overconsume nutrients and overproduce many metabolites (**Fig. 2n,o**).

### Hematopoietic stem and progenitor cells without TCA cycling increase their biosynthesis *in vivo*

Glutamine is a major carbon source for proliferating cells^35,36^. To test if loss of the TCA cycle increases glutamine utilization *in vivo,* we infused mice with U^13^C-glutamine (**Fig. 3a**). Tracer enrichment in the serum was at steady stage and did not significantly differ between *Cs^Δ/Δ^* and control mice at the time of analysis (**Extended Data Fig. 8a**). *Cs* deletion increased U^13^C-glutamine-derived labeling in bone marrow TCA cycle metabolites (**Fig. 3a**), suggesting increased activity of the linear pathway from glutamine to oxaloacetate and aspartate. *Cs* deletion also increased the contribution of glutamine to other amino acids, including alanine and proline, acetyl groups and lactate (**Fig. 3a**). Most studies of stem cell metabolism have used *in vitro* tracing^10^. To understand how nutrients contribute to *in vivo* HSPC metabolism, we performed *in vivo* stable isotope tracing for stem cells. We and others previously showed that labeling changes between treatments mostly remain stable during cell isolation^32,37^. The first turn of the TCA cycle produces M+4 citrate from U^13^C-glutamine. Conditional deletion of *Cs* abolished glutamine-derived M+4 citrate (**Fig. 3b**), consistent with a complete block of TCA cycle turning in *Cs^Δ/Δ^* HSCs and progenitors. *Cs* deletion did not induce flow of U^13^C-glutamine to M+5 citrate, consistent with oxidative and not reductive use of glutamine (**Extended data Fig. 8b**). *Cs* deletion significantly increased glutamine-derived carbon labeling in glutamate and aspartate (**Fig. 3c, d**). To test if these results reflected a generalized increase in biosynthesis, we used nitrogen-labeled glutamine. Glutamine is a major nitrogen source for the biosynthesis of amino acids and nucleotides (**Fig. 3e**). U^15^N-glutamine tracing *in vivo* revealed that *Cs* deletion elevated multiple biosynthetic pathways in HSCs or other hematopoietic cells (**Fig. 3f-k**), including glutamine-dependent synthesis of amino acids, glutathione, nucleotides, and other metabolites.

**Figure 3.**
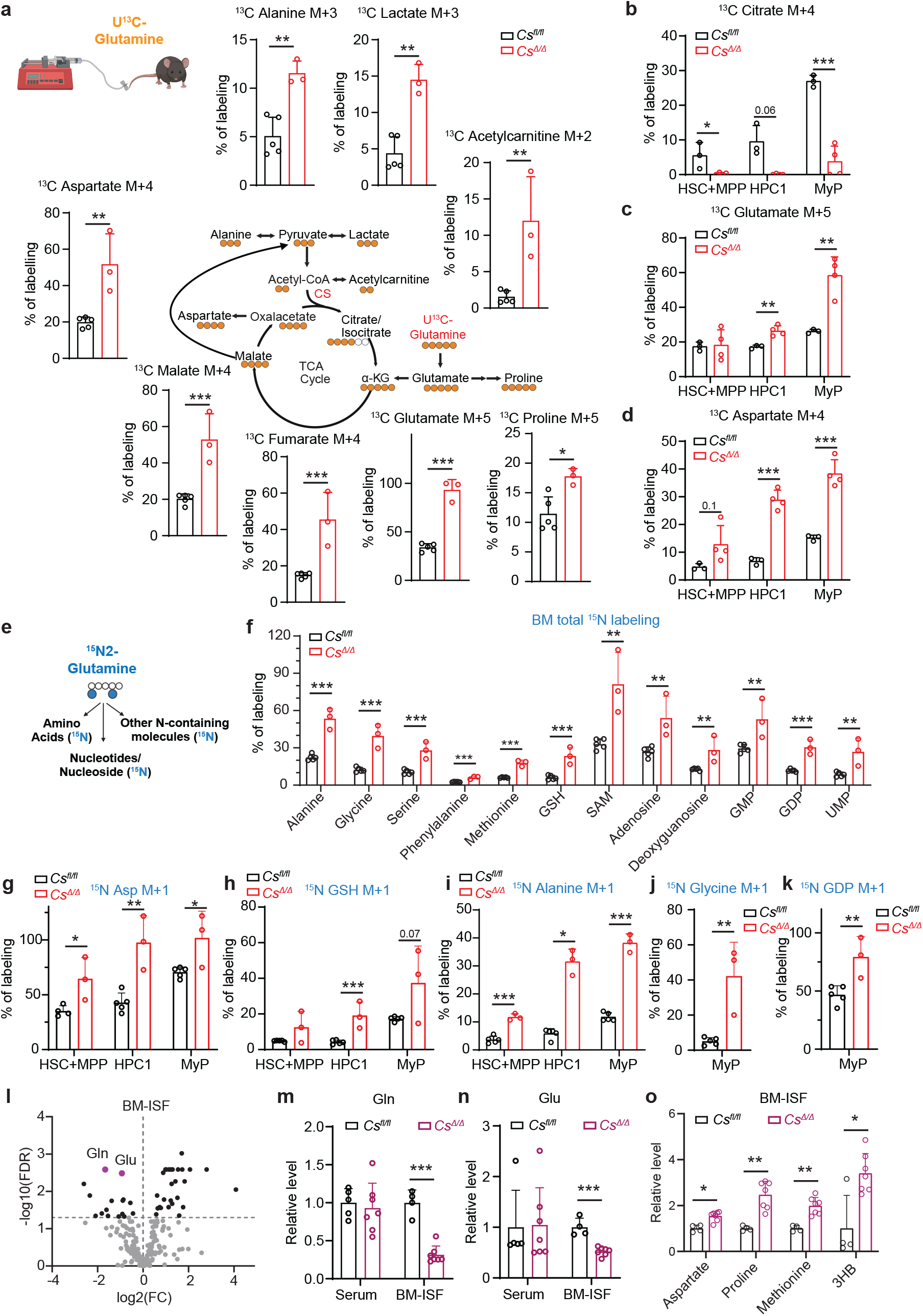
Loss of TCA cycling increases HSPC biosynthesis *in vivo*. a-d. U^13^C-glutamine *in vivo* tracing and fractional enrichment for the indicated metabolite isotopologues in BM cells (a) and in sorted HSPC populations (b-d) from *Mx1Cre;Cs^Δ/Δ^* mice or *Cs^fl/fl^* littermate controls. e-k. ^15^N2-glutamine *in vivo* tracing. Illustration of glutamine-derived nitrogen contributions (e), and fractional enrichment for the indicated metabolite isotopologues in BM cells (f), and in sorted HSPC populations (g-k). l-o. Metabolomics of the BM interstitial fluid (ISF) in *Mx1Cre;Cs^Δ/Δ^* mice or *Cs^fl/fl^* littermate controls. All data represent mean ± s.d. Statistical significance was assessed with a t-test of log-transformed data (a, f, h-HPC1, MyP, j), Mann-Whitney test (b-HPC1), Welch’s t-test (c-MyP), or with multiple t-tests of log-transformed data followed by multiple comparisons correction (l-o), and t-test (rest).

To test if the erythroid, B and T cell types that rely on *Cs* are unable to elevate glutamine use after *Cs* deletion, we infused mice with U^13^C-glutamine. *Cs* deletion arrests erythroid differentiation at the transition between Ter119^-^CD71^+^ erythroid progenitors and Ter119^+^CD71^+^ erythroblasts (**Extended Data Fig. 2k**). *Cs^Δ/Δ^* Ter119^-^CD71^+^ cells robustly increased U^13^C-glutamine-derived M+4 aspartate but this increase was mostly abrogated in *Cs^Δ/Δ^* Ter119^+^CD71^+^ cells (**Extended Data Fig. 8c)**. Therefore, erythroblast differentiation limits the ability of *Cs^Δ/Δ^* cells to use glutamine. This is consistent with the requirement of erythroblasts to synthesize rather than use glutamine to remove ammonia from heme synthesis^38^. In contrast, *Cs*^Δ/Δ^ B and T cell progenitors elevated glutamine use for aspartate synthesis, suggesting their developmental arrest was not due to limited glutamine use (**Extended Data Fig. 8d, e)**.

The microenvironment interstitial fluid (ISF) forms the direct nutrient reservoir of cells *in vivo*. We reasoned that metabolites overconsumed by *Cs^Δ/Δ^* hematopoietic cells will be depleted and those overproduced enriched in the *Cs^Δ/Δ^* ISF. Metabolomic analysis of the bone marrow ISF showed that glutamine and glutamate were the most depleted metabolites (**Fig. 3l, Supplementary Table 8**). Their reduction was specific to the ISF but not the serum (**Fig. 3m, n**), consistent with locally increased consumption by *Cs*-deficient hematopoietic cells. The levels of several amino acids (**Fig. 3o**) were elevated in the ISF, consistent with their increased intracellular levels and biosynthesis. Therefore, multiple lines of convergent evidence from *ex vivo* metabolite consumption and production analysis **(Fig. 2h-o, Extended Data Fig. 7)**, metabolomics (**Fig. 2d-g, Extended Data Fig. 4-6**), *in vivo* stable isotope tracing **(Fig. 3a-k, Extended Data Fig. 8)** and ISF metabolomics **(Fig. 3l-o)** suggest that the major metabolic role of the TCA cycle in hematopoietic stem and progenitor cells is to suppress excessive uptake of nutrients, such as glucose and glutamine, and limit biosynthesis.

### HSPCs without a turning TCA cycle outcompete HSPCs with a turning TCA cycle during regeneration

HSCs regenerate the hematopoietic system after bone marrow transplantation. We hypothesized that the increased ability of *Cs^Δ/Δ^* HSPCs to consume nutrients and divert them to biosynthesis as compared to wild-type HSPCs would increase their regenerative ability. To test this, we set up transplantation experiments in which *Cs^Δ/Δ^* and wild-type HSCs in the same environment compete to reconstitute hematopoiesis. *Mx1Cre;Cs^Δ/Δ^* or *Cs^fl/fl^*littermate control hematopoietic cells were mixed with wild-type competitors and transplanted into lethally irradiated recipient mice (**Fig. 4a**). Donor HSCs were efficiently deleted (**Extended Fig. 9a**). *Cs* deletion dramatically increased total cell and myeloid reconstitution in the blood and the marrow (**Fig. 4a-b, Extended Data Fig. 9b**). In the bone marrow, *Cs* deletion increased progenitor reconstitution (**Fig. 4c**). A second *Mx1Cre;Cs^Δ/Δ^* line from a separate genome-engineered founder mouse similarly showed increased myeloid and progenitor reconstitution (**Extended Data Fig. 9c-f).**

**Figure 4.**
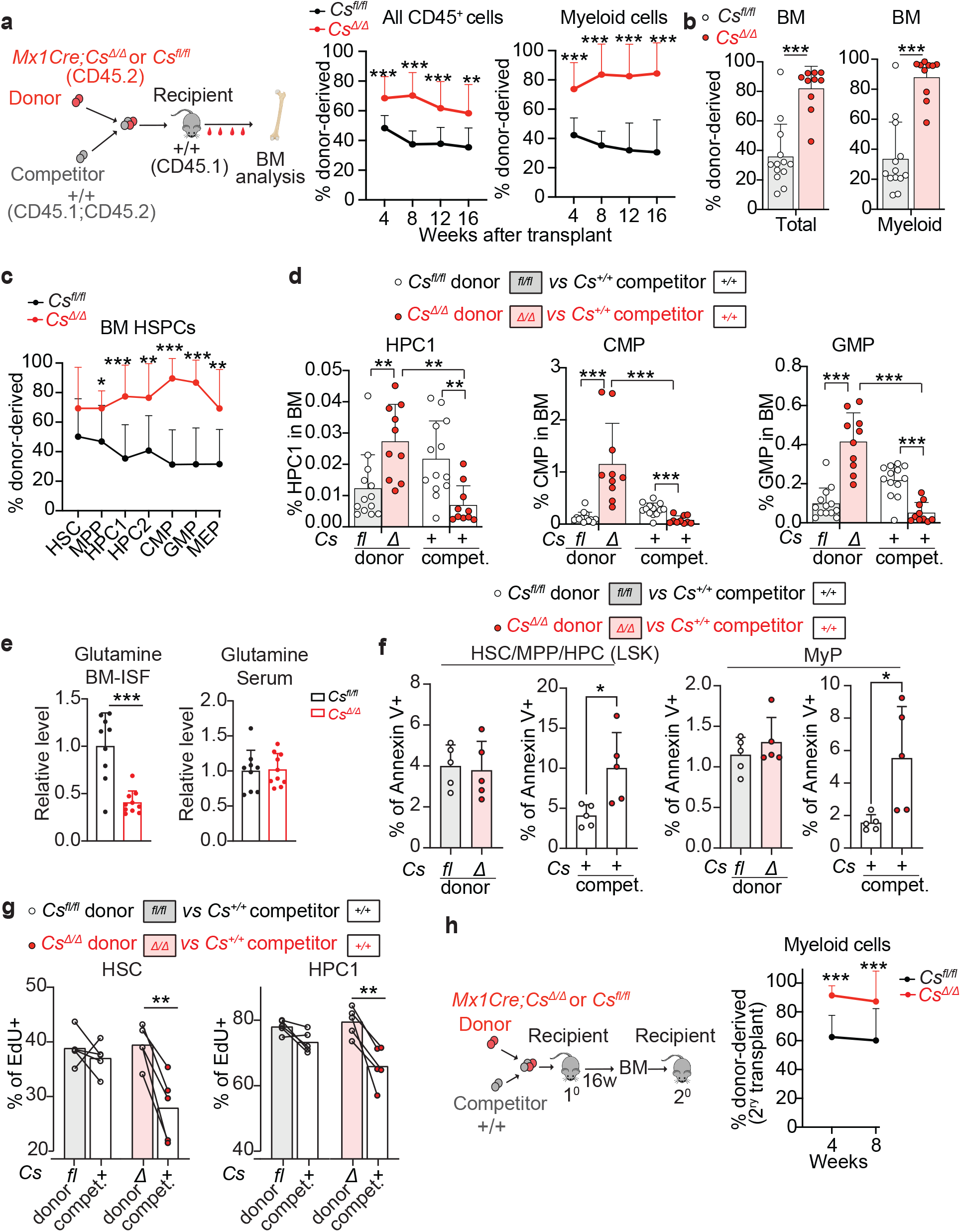
HSPCs without a turning TCA cycle outcompete HSPCs with a turning TCA cycle during regeneration. a-d. Competitive bone marrow transplantation of 500,000 donor *Mx1Cre;Cs^Δ/Δ^* or *Cs^fl/fl^* bone marrow cells with 500,000 wild-type competitor cells into lethally irradiated recipient mice. Shown are the fraction of donor-derived total CD45^+^ or myeloid cells in the blood (a) and bone marrow (b), of donor-derived HSPCs in the bone marrow (c), and the percentage of donor or competitor progenitor cells out of total bone marrow cells (d) (n=13-15 mice/genotype for blood analysis and 10-13 mice/genotype for bone marrow analysis from 3 independent experiments). e. Glutamine levels in the BM-ISF and serum of transplant recipient mice. f. Proportion of Annexin V-positive donor or competitor HSPC and myeloid progenitors in transplant recipients. g. Proportion of EdU-positive donor or competitor HSC and HPC1 after EdU administration to recipient mice for 2 days before analysis. h. Fraction of donor-derived myeloid blood cells after secondary transplants of 5 million bone marrow cells from primary transplant recipients (n=15 secondary recipients/genotype from 7 donors/genotype). All data represent mean ± s.d. Statistical significance was assessed with a t-test (a-CD45^+^ cells, c-HSC, MPP, HPC1, HPC2), Mann-Whitney test (a-Myeloid cells, b, c-CMP, GMP, MEP, d-*CS^Δ/Δ^* vs *Cs^fl/fl^* comparisons, h), Welch’s t-test (e-glutamine in ISF, g), and paired t-test (d and f-donor vs competitor comparisons).

In chimeric transplant recipients, the frequency of the *Cs^Δ/Δ^* donor HPCs, CMPs, and GMPs increased as compared to wild-type donors (**Fig. 4d**) and the frequency of the wild-type competitors of *Cs^Δ/Δ^* cells decreased as compared to wild-type competitors of *Cs^fl/fl^* cells (**Fig. 4d**). This suggested that the presence of *Cs^Δ/Δ^* cells impairs their wild-type competitors. Chimeric recipients of *Cs^Δ/Δ^* cells had depleted glutamine from the bone marrow ISF (**Fig. 4e**), similar to *Mx1Cre;Cs^Δ/Δ^* mice (**Fig. 3l-m**). *Cs^Δ/Δ^*cells impaired the proliferation and increased the apoptosis of their wild-type competitor HSCs and progenitors at homeostasis (**Fig. 4f,g**). Therefore, loss of CS cell-autonomously increases the competitiveness of HSPCs in part by impairing their wild-type competitor HSPCs. Overall, these data suggest that not only is the closed-circuit TCA cycle dispensable for stem cells, but stem cells without TCA cycle turning outcompete stem cells with TCA cycle turning.

*Cs* deletion decreased T cell reconstitution, and increased B cell reconstitution at early timepoints after transplantation, followed by a decline (**Extended Data Fig. 9g)**. *Cs* deletion increased donor-derived chimerism in B cell progenitors in the bone marrow (**Extended Data Fig. 9h)**, in line with increased production from their upstream HSPCs, however donor-derived chimerism declined at the IgM^+^ B cell stage. In the thymus, *Cs* deletion increased donor-derived chimerism of early T cell progenitors, consistent with increased marrow HSPC output, but reduced it at the double positive (DP) thymocytes (**Extended Data Fig. 9i)**. Therefore, *Cs* is cell-autonomously required in at least two distinct steps of lymphoid development.

To test if *Cs-*deficient HSCs maintained increased competitiveness after serial regenerative challenge, we performed secondary transplants. *Cs*-deficient donor cells continued to show increased myeloid reconstitution (**Fig. 4h**) and loss of T cell reconstitution (**Extended Data Fig. 9j)**. By 8 weeks post-secondary transplant, 11/15 mice with *Cs^Δ/Δ^* donor cells had >90% donor-derived myeloid reconstitution, suggesting myelopoiesis in the recipients was taken over by *Cs*-deficient cells. *Cs^Δ/Δ^* secondary recipients died at 12-16 weeks post-transplant with severe anemia, consistent with ineffective *Cs^Δ/Δ^* erythropoiesis (**Extended Data Fig. 9k**).

To avoid non-cell-autonomous effects of *Cs* deletion on HSPCs that could occur before transplant, we transplanted undeleted *Mx1Cre;Cs^fl/fl^*or *Cs^fl/fl^* bone marrow cells with wild-type competitors into lethally irradiated mice and deleted *Cs* in chimeric recipients 6 weeks later. *Cs^Δ/Δ^* HSPC and myeloid reconstitution increased, and lymphoid reconstitution decreased in the blood and bone marrow (**Extended Data Fig. 9l-o**). Similar to *Cs* deletion before transplant, the frequency of donor-derived *Cs^Δ/Δ^* HSPCs increased, and the frequency of their wild type competitors was suppressed (**Extended Data Fig. 9p**). Therefore, *Cs* deletion cell-autonomously increases HSC regeneration and myelopoiesis and impairs lymphopoiesis.

### The effects of *Cs* deletion on HSC function are not phenocopied by *Acly* deletion

In addition to fueling TCA cycle turning, CS may also provide citrate for cytosolic export and cleavage by ATP-citrate lyase (ACLY) (**Extended Data Fig. 10a**). Comparing the phenotypes of *Cs* and *Acly* deletion can dissociate *Cs* deletion phenotypes due to blocking TCA cycle turning *vs* citrate use in the cytosol. Previous work suggested that ACLY inhibition may reduce HSPCs *in vitro* or during regeneration after chemotherapy^39,40^, however, the effects of conditional *Acly* deletion on hematopoiesis have not been tested. We generated *Mx1Cre;Acly^fl/fl^* mice to do that. *Acly* was efficiently deleted in HSCs and ACLY was eliminated (**Extended Data Fig. 10b-c**). *Acly* loss caused bone marrow and spleen hypocellularity (**Extended Data Fig. 10d**). It did not affect bone marrow HSCs or restricted progenitors, increased HPCs, and had minor effects in the spleen (**Extended Data Fig. 10e-i**). *Acly* deletion decreased monocytes but not neutrophils, erythroid cells or platelets (**Extended Data Fig. 10j-n**). Most of these phenotypes differed from those of *Cs* loss. *Acly* deletion impaired bone marrow B cell development (**Extended Data Fig. 10o-p**). Most effects of *Acly* deletion persisted at 8 weeks post deletion (**Extended Data Fig. 11a-h**). By this time-point, *Acly^Δ/Δ^* mice had a smaller thymus with reduced T cell progenitors (**Extended Data Fig. 11i**). In competitive transplantations, *Acly* deletion reduced B and T lymphoid cell reconstitution (**Extended Data Fig. 11j-l**). The cell-autonomous requirements of both CS and ACLY in B and T cell development suggested a requirement for cytosolic citrate use rather than TCA cycle turning in these lineages. *Acly* deletion, in contrast to *Cs* deletion, did not affect long-term myeloid reconstitution or the proportion of donor-derived HSCs or progenitors in competitive transplantations (**Extended Data Fig. 11m**). Therefore, *Acly* deletion does not cell-autonomously affect HSC and early progenitor function, and the effects of CS loss in HSCs are not mediated by loss of ACLY-mediated cytosolic citrate use.

### The effects of *Cs* deletion on HSC function and regeneration are cell-autonomous and generalizable

To test if *Cs* deletion increases the function of purified HSCs, we transplanted highly purified EPCR^+^CD150^+^CD48^-^Lin^-^Sca-1^+^Kit^+^ HSCs with wild-type total bone marrow competitor cells to irradiated recipient mice (**Fig. 5a**). The frequency of EPCR^+^HSCs did not change in *Mx1Cre;Cs^Δ/Δ^* mice (**Extended Data Fig. 12a)**. Similar to the results of total bone marrow transplantation, *Cs* deletion conferred a large competitive advantage to EPCR^+^HSCs in blood myeloid cell and bone marrow HSPC reconstitution, along with reduced T cell reconstitution (**Fig. 5b** and **Extended Data Fig. 12b**).

**Figure 5.**
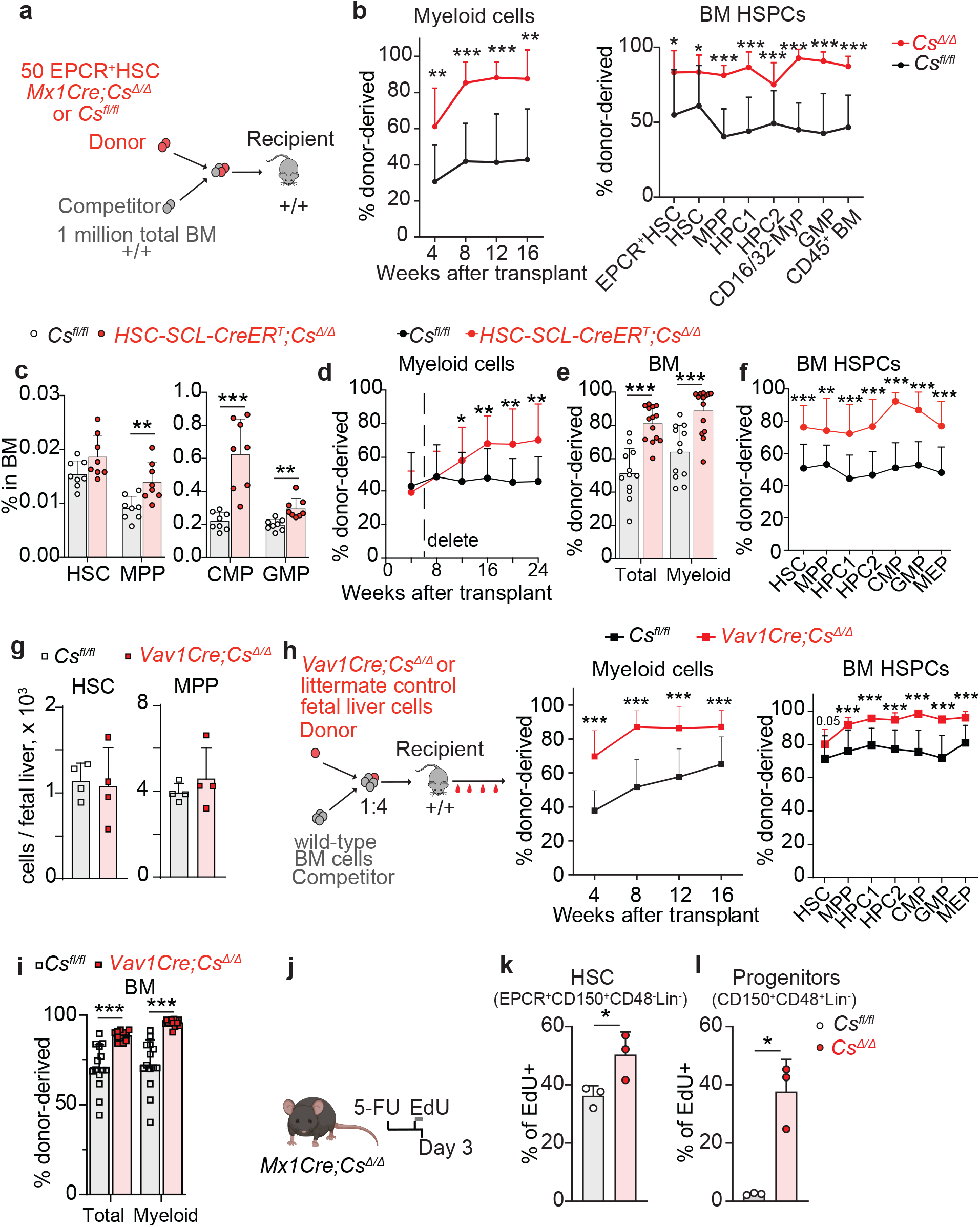
The effects of *Cs* deletion on HSC function and regeneration are cell-autonomous and generalizable. a-b. Competitive bone marrow transplantation of 50 purified *Mx1Cre;Cs^Δ/Δ^*or *Cs^fl/fl^* EPCR^+^HSCs donor cells with 1,000,000 wild-type total BM competitor cells into lethally irradiated recipient mice. Shown are the fraction of donor-derived myeloid cells in the blood (b-left) and HSPCs in the bone marrow (b-right). (n=7-12 mice/genotype from 3 experiments). c. The percentage of stem and progenitor cells in the bone marrow of *HSC-SCL-CreER^T^;Cs^fl/fl^* mice or littermate controls 3 weeks after tamoxifen administration. d-f. Competitive bone marrow transplantation of 2,000,000 donor *HSC-SCL-CreER^T^;Cs^Δ/Δ^*or *Cs^fl/fl^* bone marrow cells with 2,000,000 wild-type competitor cells into lethally irradiated recipient mice followed by tamoxifen administration 6 weeks post-transplant. Shown are the fraction of donor-derived myeloid cells in the blood (d), of donor-derived total CD45^+^ or myeloid cells in the bone marrow (e), and of donor-derived HSPCs in the bone marrow (f) (n=13-15 mice/genotype for blood analysis and 12-14 mice/genotype for bone marrow analysis from 3 independent experiments). g. Number of HSCs and MPPs in the fetal liver of E14.5 *Vav1Cre;Cs^Δ/Δ^* or littermate controls. h-i. Competitive bone marrow transplantation of 500,000 donor *Vav1Cre;Cs^Δ/Δ^*or littermate control fetal liver cells with 2,000,000 wild-type competitor total BM cells into lethally irradiated recipient mice. Shown are (h) the fraction of donor-derived myeloid cells in the blood or of donor-derived HSPCs in the bone marrow, and (i) of donor-derived total CD45^+^ or myeloid cells in the bone marrow. (n=13-14 mice/genotype from 3 independent experiments). j-l. Schematic of 5-FU administration to *Mx1Cre;Cs^Δ/Δ^* or littermate control mice followed by EdU administration for 14 hours before analysis at day 3 after 5-FU. Shown are the proportions of EdU-positive HSC and progenitor cells. All data represent mean ± s.d. Statistical significance was assessed with a t-test (b-EPCR+HSC, HSC, c-MPP, GMP, d, e, f, k), Welch’s t-test (c-CMP, h-MPP, HPC1, GMP, l), or Mann-Whitney test (rest).

To avoid the transient inflammatory effects caused by poly I:C we tested the effects of deleting *Cs* in HSPCs using *HSC-Scl-CreERT*^41^ (**Fig. 5c**). Deletion was verified by genotyping of single-HSC-derived colonies (**Extended Data Fig. 12c**). *HSC-Scl-CreER^T^;Cs^Δ/Δ^*mice had normal bone marrow cellularity 3 weeks after deletion, consistent with the absence of widespread deletion in most marrow cells^41^ (**Extended Data Fig. 12d**). *HSC-Scl-CreER^T^;Cs^Δ/Δ^*mice had preserved HSCs and HPCs and increased MPPs and some myeloid progenitor cells, similar to *Mx1Cre;Cs^fl/fl^*mice (**Fig. 5c, Extended Data Fig. 12e-g**). Τransplantation of undeleted *HSC-Scl-CreER^T^;Cs^fl/fl^* bone marrow cells followed by tamoxifen-mediated deletion 6 weeks later, showed increased myeloid and HSPC reconstitution (**Fig. 5d-f, Extended Data Fig. 12h,i**), phenocopying *Mx1Cre;Cs^fl/fl^* mice. The frequency of donor-derived *Cs^Δ/Δ^* HSPCs increased, and the frequency of their wild-type competitors was suppressed (**Extended Data Fig. 12j**). Therefore, *Cs* deletion cell-autonomously increases HSC function and their ability to outcompete wild-type HSCs.

Fetal liver HSCs are more proliferative and have increased long-term reconstitution potential compared to adult HSCs^42^. To test if loss of TCA cycle turning also promotes HSC function in this potent stem cell population, we generated *Vav1Cre;Cs^fl/fl^* mice to delete *Cs* in fetal hematopoiesis. No *Vav1Cre;Cs^Δ/Δ^* mice were born, suggesting embryonic lethality (**Extended Data Fig. 13a**). We therefore analyzed fetal liver hematopoiesis at E14.5. *Vav1Cre;Cs^Δ/Δ^* were pale (**Extended Data Fig. 13b**), indicating anemia, and erythroid differentiation was arrested at the Ter119^+^CD71^+^ erythroblast stage (**Extended Data Fig. 13c**), consistent with effects of *Cs* deletion in the adult. *Cs* deletion did not reduce the number of other hematopoietic cell types, including HSCs, progenitors, or myeloid cells, suggesting that the TCA cycle is dispensable for fetal HSPCs (**Fig. 5g**, **Extended Data Fig. 13d)**. Transplantation of E14.5 *Vav1Cre;Cs^Δ/Δ^* or littermate control fetal liver cells with wild-type total bone marrow competitor cells showed that *Cs* deletion increased myeloid and HSPC reconstitution, similar to the adult *Cs* knockout models we analyzed (**Fig. 5h-i, Extended Data Fig. 13e**). Therefore, loss of TCA cycle turning promotes the function of fetal HSCs similar to its effects in adult HSCs.

To test if loss of TCA cycle turning promotes regeneration in an orthogonal model, we used 5-fluorouracil (5-FU) challenge, which induces HSCs to proliferate and replenish the ablated hematopoietic compartment. The kinetics of HSC function in this model are well characterized, with the onset of HSC proliferation at day 3 and most HSCs proliferating by day 4^43^. To test whether *Cs* deletion accelerates the onset of proliferation, we administered EdU to *Mx1Cre;Cs^Δ/Δ^*mice with 2 injections 2.5 days after 5-FU treatment and analyzed EdU incorporation on day 3 (**Fig. 5j**). *Cs* deletion increased the proportion of EdU-positive EPCR^+^CD150^+^CD48^-^Lin^-^ HSCs, defined with surface markers for functional HSCs after 5-FU^43–45^ (**Fig. 5k**). *Cs* deletion increased the proportion of EdU-positive Lin^-^CD150^+^CD48^+^ hematopoietic progenitor cells (**Fig. 5l**), which are among the first cells produced from proliferating HSCs after challenge^40,45^. *Cs* deletion increased the frequency of HSCs and progenitors at this early time point (**Extended Data Fig. 13f)**, consistent with accelerated proliferation. To test regeneration of hematopoiesis, we used *HSC-SCL-CreERT;Cs^Δ/Δ^*mice to avoid the impaired erythroid and lymphoid development of *Mx1Cre;Cs^Δ/Δ^*mice. *Cs* deletion accelerated blood neutrophil recovery (**Extended Data Fig. 13g)** and increased the number of multipotent and myeloid progenitors in the bone marrow (**Extended Data Fig. 13h)**. Therefore, blocking TCA cycle turning accelerates the onset of HSC proliferation and increases production of their progeny during regeneration.

### Loss of TCA cycle turning increases HSC function by increasing nutrient use and biosynthesis

To understand if changes in gene expression levels underpinned nutrient overconsumption and biosynthesis caused by *Cs* deletion, we performed RNA-seq analysis of *Mx1Cre;Cs^Δ/Δ^* or littermate control CD48^-^Sca-1^+^Kit^+^Lin^-^ HSCs + MPPs. Expression of TCA cycle genes other than *Cs* did not significantly change, consistent with the idea that *Cs* deletion specifically disrupted cycle turning (**Extended Data Fig. 14a**). 31 genes were differentially expressed at an FDR level < 0.05 (**Fig. 6a, Supplementary Table 9**). Slc2a1 (GLUT1), which imports glucose in HSCs^46^, was upregulated (**Extended Data Fig. 14b**). Three of the top four differentially expressed genes were canonical targets of the ATP-responsive transcription factor MondoA, including *Txnip* and *Arrdc4*, which were downregulated in *Cs^Δ/Δ^* HSCs (**Fig. 6a,b)**. *Txnip* and *Arrdc4* are elevated in response to high mitochondrial ATP to limit glucose uptake by removing cell surface GLUT1 through endocytosis^47–49^. Their downregulation was consistent with the reduced ATP levels of *Cs^Δ/Δ^* HSCs (**Fig. 2b**). *Cs^Δ/Δ^* HSCs and other progenitors showed a 6-fold increase in GLUT1 surface levels, explaining their increased glucose uptake (**Fig. 6c**). Given the increased glutamine uptake of *Cs^Δ/Δ^* HSPCs, we assessed glutamine transporter levels. ASCT2 and SNAT2 levels were 2-4 fold elevated in *Cs^Δ/Δ^* HSPCs (**Fig. 6d, Extended Data Fig. 14c**). Therefore, *Cs* deletion elevates glucose and glutamine transporters to increase nutrient consumption. In addition to decreased ATP leading to *Txnip/Arrdc4* repression, *Cs* deletion may affect transcription through other mechanisms, such as impairing the activity of TET enzymes. Levels of the TET product 5-hydroxymethyl-2-deoxycytidine (5hmC) were reduced in the DNA of *Cs^Δ/Δ^*HSCs and progenitors (**Extended Data Fig. 14d**), suggesting reduced TET activity.

**Figure 6.**
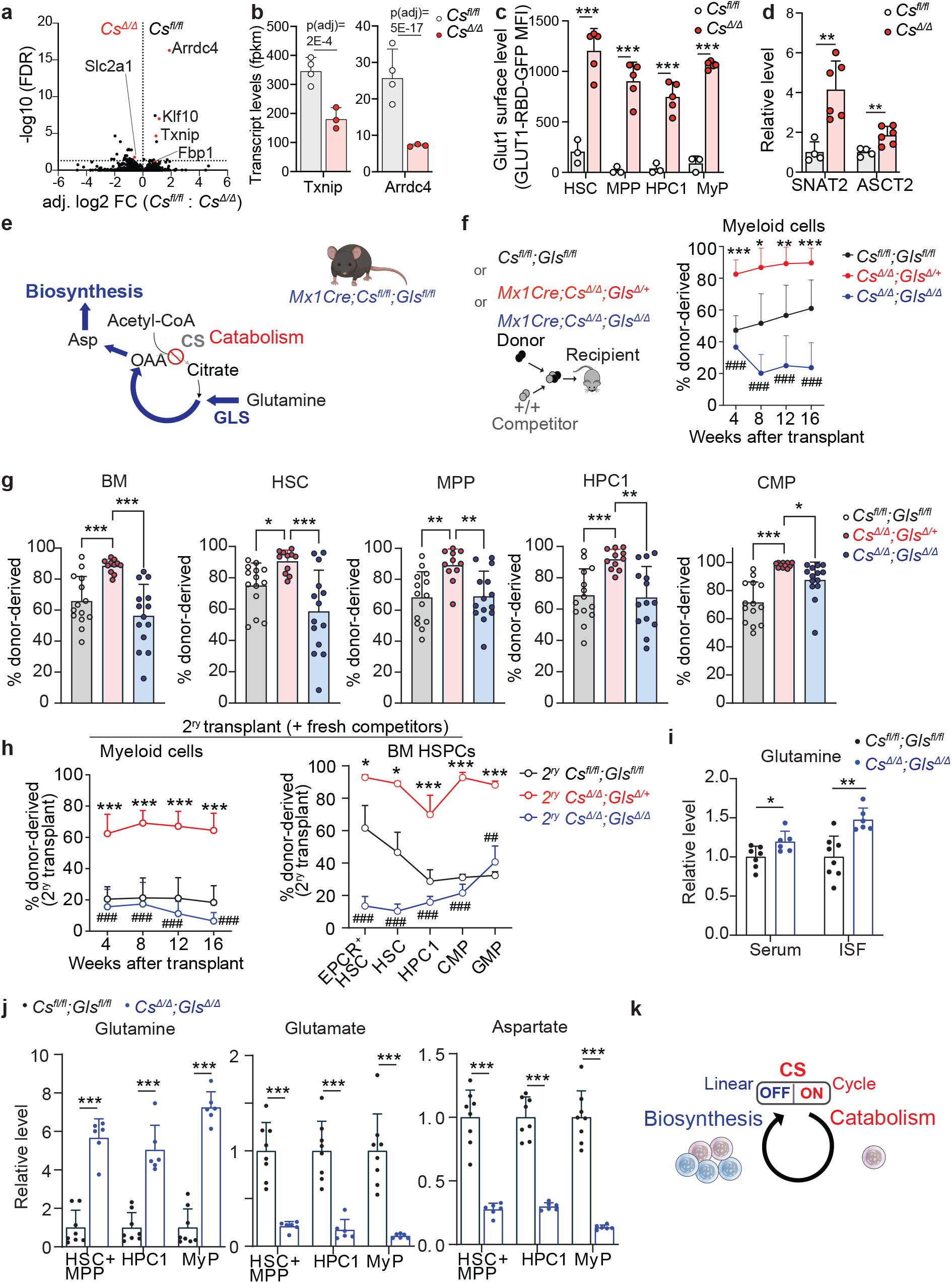
Loss of TCA cycle turning increases HSC function by increasing nutrient use and biosynthesis. a-b. Volcano plot showing differential gene expression from RNA-seq data from double-sorted CD48^-^Lin^-^ Sca-1^+^Kit^+^ HSC+MPP from *Mx1Cre;Cs^Δ/Δ^* mice or *Cs^fl/fl^* littermate controls (n=3-4 mice/genotype) and expression levels of Txnip and Arrdc4. The dotted horizontal line in the volcano plot represents an FDR < 0.05 cutoff. c. GLUT1 surface levels in indicated HSPC populations from *Mx1Cre;Cs^Δ/Δ^*or *Cs^fl/fl^* mice measured by GLUT1.RBD median fluorescence intensity over background using flow cytometry d. SNAT2 and ASCT2 levels in sorted Lin^-^Kit^+^ HSPCs from *Mx1Cre;Cs^Δ/Δ^*or *Cs^fl/fl^* mice measured using a western blot. e. Schematic showing that CS loss blocks full nutrient catabolism and increases glutamine-mediated biosynthesis and use of the *Mx1Cre;Cs^Δ/Δ^;Gls^Δ/Δ^* mouse to test this. f-g. Competitive bone marrow transplantation of 2 million donor *Mx1Cre;Cs^Δ/Δ^;Gls^Δ/Δ^*or *Mx1Cre;Cs^Δ/Δ^;Gls^Δ/+^* or *Cs^fl/fl^Gls^fl/fl^* bone marrow cells with 2 million wild-type competitor cells into lethally irradiated recipient mice. Shown are (f) the fraction of donor-derived myeloid cells in the blood and (g) donor-derived HSPCs in the bone marrow (n=14-15 mice/genotype for blood analysis and 11-14 mice/genotype for bone marrow analysis from 3 independent experiments). h. Fraction of donor-derived myeloid blood cells and donor-derived HSPCs in the bone marrow after secondary transplants of 5 million bone marrow cells from primary transplant recipients plus 5 million fresh competitor bone marrow cells (n=13-15 mice/genotype for blood analysis and 3-4 mice/genotype for bone marrow analysis from 3 independent primary donors). i-j. Glutamine levels in the BM ISF and serum (i) or metabolite levels in sorted HSC and progenitor populations (j) of *Mx1Cre;Cs^Δ/Δ^;Gls^Δ/Δ^*or littermate control mice. k. Illustration showing the role of CS in regulating biosynthesis. All data represent mean ± s.d. Statistical significance was assessed with a one-way ANOVA (f-4w, 16w, g-MPP, h-HPC1), Kruskal-Wallis test (f-8w, 12w, g-HSC, CMP, h-myeloid cells), Welch one-way ANOVA (g-BM, HPC1, h-EPCR+HSC, HSC, CMP, GMP), or Welch’s t-test (c, j-glutamate-HSC+MPP, MyP, Aspartate), Mann-Whitney t-test (j-glutamine-HSC+MPP, j-glutamate-HPC1), and t-test (d, i). RNA-seq statistical analysis was performed with DESeq2 followed by the Benjamini-Hochberg method to control the false discovery rate as described in the methods.

To understand if metabolic or epigenetic effects were required for the effects of *Cs* deletion on HSC function, we generated *Mx1Cre;Cs^fl/fl^;Gls^fl/fl^*mice (**Fig. 6e**). Glutaminase (GLS) converts glutamine to glutamate for use in oxidative and biosynthetic reactions. *Gls* deletion does not affect HSC function in competitive transplantations^50^. Transplantation of *Mx1Cre;Cs^Δ/Δ^;Gls^Δ/Δ^*bone marrow cells in competition with wild-type cells showed that *Gls* deletion blocked the increased *Cs^Δ/Δ^* HSC function as measured by peripheral blood and HSPC bone marrow chimerism (**Fig. 6f-g**). To test if *Cs^Δ/Δ^* HSPCs maintained their advantage in serial transplants beyond 8 weeks (**Fig. 4h**, **Extended Data Fig. 9k)** and if this was abrogated by *Gls* deletion, we performed secondary transplants with fresh wild-type competitors. *Cs^Δ/Δ^*cells maintained a large competitive advantage for at least 16 weeks in the blood and 6 months in the HSPC compartment of secondary recipients, and this was abolished by *Gls* deletion (**Fig. 6h**). We then tested if the rescue of increased *Cs^Δ/Δ^* HSC function by *Gls* deletion was due to a rescue of metabolic or epigenetic effects of *Cs* deletion. *Cs^Δ/Δ^;Gls^Δ/Δ^* HSPCs had an even stronger 5hmC loss than *Cs^Δ/Δ^* HSPCs, suggesting *Gls* deletion did not rescue 5hmC loss caused by *Cs* deletion and arguing against the idea that 5hmC loss promoted *Cs^Δ/Δ^* HSC function (**Extended Data Fig. 14e**). In contrast, *Gls* deletion rescued the depletion of glutamine in the BM-ISF of *Cs^Δ/Δ^* mice (**Fig. 6i**), indicating that glutamine ISF depletion was caused by increased glutamine use. *Cs^Δ/Δ^;Gls^Δ/Δ^*HSCs and progenitors had severely reduced glutamate and aspartate and increased glutamine levels, suggesting *Gls* deletion blocked the biosynthetic pathway from glutamine to aspartate in *Cs^Δ/Δ^* HSPCs (**Fig. 6j**). Therefore, increased glutamine use in this pathway is required for the increased stem cell function caused by loss of TCA cycle turning.

## Discussion

The TCA cycle serves as the universal terminal step in the complete oxidation of all nutrients to sustain respiration and energy production^1^. The cycle is also thought to contribute to biosynthesis^3,6,7^. However, the *in vivo* necessity of cycle turning itself had not been tested. CS delivers the nutrient fuel oxidized in the cycle. By conditionally deleting *Cs* in a somatic stem cell system, we found that, surprisingly, TCA cycle turning was dispensable *in vivo* for respiration, cell survival, anabolism, proliferation, and function of hematopoietic stem and progenitor cells. CS was required at specific steps of lymphoid and erythroid development for reasons other than to support full nutrient combustion. *Cs* germline mutations in flies are lethal when null or cause photoreceptor degeneration when hypomorphic^51,52^. Thus, CS is required in some specialized cell types, but TCA cycle turning is not generally required for the major metabolic roles classically ascribed to it.

A major goal of regenerative medicine is to identify manipulations that can enhance HSC function and hematopoietic regeneration. This is not a trivial problem because at baseline HSCs are defined by their extraordinary regenerative ability^53^ and are the most fecund somatic stem cells in the body^13,14^. Reducing CS activity provides a method to accomplish this. Aspects of mitochondrial metabolism are dispensable for HSCs. For example, PDH deletion, which blocks glucose oxidation but, unlike *Cs* deletion, does not block oxidation of all nutrients in the cycle, does not change HSC function^37^, and conversely, PDH activation inhibits HSC function^19^. However, several mitochondrial proteins are required for HSC function^10,23–25,34,54^. Our work shows that it is complete nutrient oxidation in the cyclic pathway, rather than oxidative metabolism per se that limits HSC function. *Cs-*deficiency caused HSPCs to overconsume nutrients, increase biosynthesis, and deplete the microenvironment from glutamine to outcompete wild-type HSCs. *Cs* deletion increased glutamine use, which promotes proliferation by providing nitrogen and carbon for nucleotide and amino acid synthesis^36,55,56^, and this was required for the effects of *Cs* deletion on HSCs. One of the main glutamine outputs in *Cs*-deficient HSCs was aspartate. Elevated aspartate promotes HSC function by stimulating protein and nucleotide synthesis^34^ and promotes proliferation of cancer cells^33,57^. Therefore, *Cs* loss increases HSC function and regenerative ability in part by increasing glutamine use and biosynthesis (**Fig. 6k**). *Cs* loss causes additional molecular changes that promote HSC function and regeneration, such as enhancing glucose uptake, and depleting nutrients from the microenvironment to increase competition. Thus, CS coordinates multiple aspects of metabolism to regulate stem cell function.

Cancer cells tend to have high nutrient uptake, glycolysis, glutaminolysis, and lactate production, and low TCA cycle activity rates as compared to normal cells^11,12,58,59^. These metabolic adaptations are similar to those we observe in *Cs-*deficient HSPCs. CS competes with GOT2 for oxaloacetate. Funneling of nutrients to terminal oxidation by CS diverts oxaloacetate from aspartate synthesis. Therefore, at the fundamental metabolic level, the turning of the TCA cycle is antagonistic to anabolism. Consistent with this idea, partial inhibition of CS in glioblastoma cells *in vitro* increases aspartate and asparagine synthesis^35^. Our results suggest that a major function of full nutrient combustion in the TCA cycle is to reduce the amount of nutrients necessary to serve cellular needs. Thus, an intact, turning TCA cycle limits nutrient overconsumption, anabolism, and cell competition. These findings may explain why TCA cycle activity is reduced in many proliferating cells in embryonic development, adult tissues, and cancers^8–12,58,59^.

## Methods

### Mice

All mice were on a C57BL background. *Cs^fl^* mice were generated by the Children’s Research Institute Genome Engineering Core. *Cs* exon 10 was targeted because it codes for part of the CS enzymatic active site and is flanked by introns of sufficient length containing sequences suitable for CRISPR targeting. The length of most earlier *Cs* exons is short and a multiple of 3, complicating their targeting in conditional knockout strategies. A pre-assembled ribonucleoprotein complex consisting of Alt-R S.p. Cas9 Nuclease V3 and sgRNA with a single-strand DNA donor oligo was microinjected into C57BL/6J zygotes. Targeting sequences were, for the left gRNA: ACTGTACATCAAAGTCAATC, and for the right gRNA: GTGAGACCTGTCTCAACTAG. PCR analysis and sequencing were used to genotype the targeted locus in chimeric mice. 2 founder mice were each backcrossed 4 times to the C57BL/Ka-Thy-1.1 background before crossing to other lines. *Cs^fl^* genotyping was performed with PCR using primers GCATTTTATGGCTAGGCAAG, GGTCAGTCTTCCTCAGTACTG, GAGTGGCATCAATGACCTCC which amplified bands of 249 bp for the wild-type allele, 289 bp for the flox allele, and 417 bp for the deleted allele. *Mx1-Cre*^60^, *HSC-Scl-CreER^T^* ^41^, and *Vav1Cre*^61^ mice were obtained from the S. Morrison lab. *Gls^fl^* mice^62^ were obtained from The Jackson Laboratory (017894). *Acly^fl^* mice^63^ were obtained from MMRRC (043555-JAX). *Cs^fl^, Gls^fl^*, and *Acly^fl^* mice were each backcrossed 4 times to the C57BL/Ka-Thy-1.1 background. Experiments were performed with both male and female mice. Mice with *Mx1Cre* and their littermate controls were injected intraperitoneally five times with 40 μg poly I:C in PBS every other day at 6-9 weeks of age. Mice with *HSC-SCL-CreER^T^* and their littermate controls were injected intraperitoneally five times with 2 mg of tamoxifen in corn oil daily at 6-9 weeks of age. Mice were analyzed at 3-4 weeks after the first poly I:C injection unless otherwise noted in the text. C57BL/Ka-Thy-1.1/Thy-1.2 (CD45.2/CD45.1) mice were used as competitors and C57BL/Ka-Thy-1.2 (CD45.1) mice as recipients in transplantation experiments. Mice were housed at the Animal Resource Center at the University of Texas Southwestern Medical Center. All procedures were approved by the UT Southwestern Institutional Animal Care and Use Committee.

### Hematopoietic analysis

Bone marrow cells were obtained by flushing femurs with a 25G needle (for analysis) or by crushing all bones using a mortar and pestle (for cell sorting or for analysis of transplant recipients) in Hank’s Balanced Salt Solution without calcium and magnesium (HBSS) + 2% heat inactivated bovine serum (HIBS). Spleens and thymuses were mechanically dissociated by crushing and trituration. Cell suspensions were filtered through a 40 μm strainer and the cell number was counted with a Vi-cell XR analyzer (Beckman Coulter). Cells were stained with fluorochrome-conjugated surface antibodies for 30 minutes at 4 °C protected from light or when CD34 antibody was included for 90 minutes on ice. After staining, cells were washed and resuspended in HBSS + 2%HIBS + 4′,6-diamidino-2-phenylindole (DAPI, 1 μg/ml) or propidium iodide (1 μg/ml) for live/dead cell identification. Hematopoietic and blood cell populations were defined with the following markers: CD150^+^CD48^-^Lineage^-^Sca-1^+^Kit^+^ hematopoietic stem cells (HSCs), EPCR^+^CD150^+^CD48^-^Lineage^-^Sca-1^+^Kit^+^ (EPCR^+^HSCs), CD150^-^ CD48^-^Lineage^-^Sca-1^+^Kit^+^ multipotent progenitor cells (MPPs), CD150^-^CD48^+^Lineage^−^Sca-1^+^Kit^+^ (HPC1), CD150^+^CD48^+^Lineage^−^Sca-1^+^Kit^+^ (HPC2), CD34^+^CD16/32^-^Lineage^-^Sca-1^-^Kit^+^ common myeloid progenitors (CMPs), CD34^+^CD16/32^+^Lineage^-^Sca-1^-^Kit^+^ granulocyte-monocyte progenitors (GMPs), CD34^-^CD16/32^-^Lineage^-^Sca-1^-^Kit^+^ megakaryocyte-erythroid progenitors (MEPs), Lineage^-^Sca-1^-^ Kit^+^CD150^+^CD41^+^ megakaryocyte progenitors (MkP), Lineage^-^Sca-1^-^Kit^+^CD41^-^CD16/32^-^CD150^+^CD105^-^ pre-megakaryocytic-erythroid progenitors (preMegE), Lineage^-^Sca-1^-^Kit^+^CD41^-^CD16/32^-^CD150^+^CD105^+^ pre-colony forming unit-erythroid (preCFU-E), Lineage^-^Sca-1^-^Kit^+^CD41^-^CD16/32^-^CD150^-^CD105^+^ colony forming unit-erythroid (CFU-E), Lineage^-^Sca-1^-^Kit^+^CD41^-^CD16/32^-^CD150^-^CD105^-^ pre-granulocyte-macrophage progenitors (preGM), Mac-1^+^ myeloid cells, Mac-1^+^Ly-6g^+^CD115^-^ neutrophils, Mac-1^+^Ly-6g^-^ CD115^+^Ly-6c^+^ inflammatory monocytes, CD71^mid^Ter-119^-^, CD71^+^Ter-119^-/low^, CD71^+^Ter-119^+^, CD71^mid^Ter-119^+^ erythroid progenitors, CD3^+^ T cells, B220^+^ B cells, Mac1^-^B220^+^IgM^-^CD43^+^CD24^-^ pre-pro-B cells, Mac1^-^B220^+^IgM^-^CD43^+^CD24^+^ pro-B cells, Mac1^-^B220^+^IgM^-^CD43^-^CD24^+^ pre-B cells, IgM^+^B220^+^ mature B cells. The Lineage antibody cocktail for HSPCs consisted of CD2, CD3, CD5, CD8, Ter-119, B220 and Gr-1 antibodies. T cell progenitor populations in the thymus were defined with the following markers, after excluding Mac-1^+^, Ter-119^+^ or B220^+^ cells: CD3^+^CD4^+^CD8^-^ (CD4^+^ single positive, SP), CD3^+^CD4^-^ CD8^+^ (CD8^+^ SP), CD3^-^CD4^-^CD8^+^ (ISPs), CD3^-^CD4^+^CD8^+^ (CD3^-^DPs), CD3^+^CD4^+^CD8^+^ (CD3^+^DPs), CD4^-^ CD8^-^ (DNs), CD4^-^CD8^-^CD44^+^CD25^-^ (DN1), CD4^-^CD8^-^CD44^+^CD25^+^ (DN2), CD4^-^CD8^-^CD44^-^ CD25^+^ (DN3), CD4^-^CD8^-^CD44^-^CD25^-^ (DN4). For 5-FU experiments, HSCs were defined as EPCR^+^CD150^+^CD48^-^Lin^-^ cells and HPC-2 progenitors as CD150^+^CD48^+^Lin^-^ cells. Antibodies were purchased from Biolegend, BD, or Tonbo. Cell samples were analyzed with a FACSCanto or LSRFortessa or FACSAria (BD). Flow cytometry data were analyzed with FlowJo (FlowJo LLC). Genotyping from single HSC-derived colonies was performed by sorting single HSCs into a 96-well flat-bottom plate containing 100 μl of Methocult (M3434, StemCell Technologies) followed 2 weeks later by DNA extraction from the resulting colonies and PCR.

### Bone marrow reconstitution assays

These were performed according to previous work^26,37^. For competitive transplants, in most experiments 500,000 donor (CD45.2) and 500,000 competitor (CD45.1/CD45.2) cells were mixed and transplanted into the retro-orbital venous sinus of anesthetized CD45.1 recipients that had been irradiated using an XRAD 320 X-ray irradiator (Precision X-Ray) with two doses of 540 rad (1080 rad in total) delivered at least 3h apart. In some experiments 2,000,000 donor + 2,000,000 competitor cells were used. As indicated in the text, donor cells were obtained from experimental or littermate control donor mice 3 weeks after the start of poly I:C administration, or in some cases before poly I:C or tamoxifen administration. In experiments in which deletion was induced after transplant, poly I:C or tamoxifen was administered at 5-6 weeks post-transplant. Recipient mice were maintained on antibiotic water (Baytril 0.08 mg/ml) for 1 week before and 4 weeks after transplantation. Peripheral blood was obtained from the tail veins of recipient mice every four weeks. Red blood cells were lysed with ammonium chloride potassium buffer. Samples were stained with CD45.2 (104), CD45.1 (A20), B220, Mac1, CD3, CD4, CD8, Ly-6c, Ly-6g, and CD115 antibodies, and analyzed with flow cytometry. At the experimental endpoint, recipient mice were euthanized, and bone marrow was obtained from crushed femurs, tibias, pelvic bones, and spine for surface antibody staining and flow cytometry analysis or for secondary transplant.

### EdU incorporation analysis

For proliferation analysis, primary mice were injected intraperitoneally with 5-ethynyl-2′-deoxyuridine (EdU, 100 mg/kg of body mass) and euthanized 2 hours later. After cell surface staining, cells were fixed in 1% formaldehyde on ice for 20 minutes, washed in 1x DPBS + 1% BSA, permeabilized with 0.1% saponin in 1x DPBS + 1% BSA for 15 minutes on ice and stained for 20 minutes at room temperature with 1x DPBS + 1 mM CuSO_4_ + 5 μΜ Azide-Alexa Fluor 555 (Thermo) + 100 mM freshly made ascorbate. Cells were washed and analyzed with flow cytometry. For proliferation analysis in transplant recipients, mice were injected intraperitoneally with EdU 3 times at 48 h, 24 h, and 2 h before euthanasia. The spine, leg and pelvic bones of each mouse were dissected and crushed for bone marrow extraction. After Kit^+^ cell magnetic enrichment and staining with cell surface antibodies, donor (104^+^A20^-^) or competitor (104^+^A20^+^) CD150^+^CD48^-^Lineage^-^Sca-1^+^Kit^+^ HSCs or CD150^-^CD48^+^Lineage^-^Sca-1^+^Kit^+^ hematopoietic progenitor cells were sorted into tubes containing 2 million unstained carrier bone marrow cells, fixed, washed, and stained as above, except 2.5 μΜ Azide-Alexa Fluor 555 was used.

### Apoptosis analysis

1 million Kit^+^ enriched cells from transplant recipients obtained as above were stained with cell surface antibodies and washed 3 times with cold DPBS before staining with APC Annexin V (BD) at RT in the dark in 200 μL 1x binding buffer for 15 minutes. 200 μL 1x binding buffer was then added to each sample before flow cytometry analysis.

### 5-FU assay

Mice were injected intraperitoneally with 5-fluorouracil (5-FU, Sigma-Aldrich, 150 mg/kg) and euthanized for analysis at day 3 or day 14. For mice at the day 3 timepoint, 2 doses of EdU were injected 14h and 2h before euthanasia, respectively, to assess the proliferation of stem and progenitor cells. Flow cytometry analysis of BM cell populations was performed with cell surface markers described in the text. EdU analysis was performed with Kit^+^ cell enrichment, sorting of EPCR^+^CD150^+^CD48^-^Lineage^-^ HSCs or CD150^+^CD48^+^Lineage^-^ hematopoietic progenitor cells into tubes containing unstained carrier cells, and click staining as described above.

### Cell surface Glut1 measurement

Glut1 binding reagent from Metafora (GLUT1.RBD) was used to measure cell-surface GLUT1 levels. BM cells were collected from all bones and Kit^+^ cells were magnetically enriched with LS columns (Miltenyi). 1 million Kit^+^ cells were stained with cell surface markers, washed twice with DPBS + 5% HIBS, and stained with 0.6 μg GLUT1.RBD reagent at 37°C for 20 minutes. Samples were washed twice with DPBS + 5% HIBS and reconstituted in DPBS + 5% HIBS with PI as a live/dead marker for flow analysis. The signal median fluorescence intensity (MFI) of each sample was calculated as the MFI for experimental samples minus the MFI of the fluorescence-minus-one control tube (FMO-stained with surface antibodies and without Glut1.RBD).

### Flow cytometry analysis of fluorescent sensors

BM cells from a femur were collected in staining media and counted. A total of 5 million cells were used for staining with the following antibody cocktails: For HSPC mitochondria membrane potential measurement with TMRM: CD150-BV421, CD48-AF700, Kit-APC750, Sca-1-PECy7, CD34-Bio, CD16/32-BV510, and lineage-FITC (including CD2, CD3, CD5, CD8, B220, Ter119, Gr1). For HSPC mitochondria mass measurement with MitoTracker Green, the antibodies were the same as above, except lineage antibodies were PE conjugated. For HSPC ROS analysis with Enzo Total ROS, the antibodies were the same as above except for CD16/32-PE and Lineage-APC conjugates. Samples were stained with surface markers on ice for 90 minutes protected from light, with mixing every 30 minutes, and then washed twice with HBSS + 0.2% BSA to remove the serum. The supernatant was removed after every wash by tapping the tube. Then, cells were resuspended in 1 ml of DMEM (Sigma D5030) pH=7.4, with added 2 mM glutamine, 1 mM pyruvate, 10 mM glucose, 50 μM verapamil to block efflux pumps, and streptavidin conjugated to APC (for TMRM and MitoTracker experiments) or to PERCPCy5.5 (for Enzo Total ROS experiments). One of the following sensors was added to each sample: TMRM 20 nM (Therrmo), MitoTracker Green 100nM (Thermo), and Enzo Total ROS Green 5 μM (Enzo). Cells were stained in a 37 °C water bath protected from light for 15 minutes, washed with HBSS + 2% HIBS, the washing buffer was completely removed, and cells were resuspended in HBSS + 2% HIBS with 50 μM verapamil and live/dead dye (PI, 1:1000). The samples were kept on ice away from light during flow cytometry analysis. The median fluorescence intensity (MFI) of each sample was normalized to the average MFI of the live cell population of the wild-type samples in each experiment.

### Cell sorting for Seahorse analysis

Femurs, tibias, pelvic bones, and spine were collected, cleaned thoroughly, crushed in 15 mL staining media (HBSS with 2%HIBS), and filtered through a 40 μm strainer. 0.1 mL of the cell suspension was set aside for total BM measurements, and the remaining cells were washed and resuspended in 1 mL staining media. Each sample was stained with 10 μL of Kit-APC780 antibody for 30 minutes at 4°C. Samples were washed and resuspended in 500 μL staining media containing 20 μL of anti-APC magnetic beads for 30 min at 4°C, protected from light. Cells were mixed halfway through each staining period, then washed, resuspended in 2 mL staining media, and filtered. Positive magnetic selection of Kit^+^ cells was performed using LS magnetic columns and a QuadroMACS manual separator (Miltenyi) in the cold room. After two on-column washes with 3 mL staining media, positively selected cells were eluted with 2 mL staining media, centrifuged, resuspended in 200 μL staining media with cell surface antibodies, stained for 90 minutes at 4°C, washed, resuspended in 800 μL staining media with DAPI (1:1000) and filtered before sorting. BD FACSymphony S6 was used to sort a total of 400,000-700,000 cells Lin^-^Kit^+^ HSPCs per mouse with a 70 μm nozzle in four-way purity sort mode to minimize the volume of sorted drops from each mouse into 1.5 mL tubes containing 100 μL of DPBS. For total BM analysis, cells were treated with ACK lysis buffer to lyse red blood cells, washed, resuspended in 1 mL of staining media, and 400,000 cells were counted using Vi-Cell XR and used for analysis.

### Seahorse Assay

The assay was done by following the Agilent Seahorse XF Cell Mito Stress Test protocol, with some modifications. The cell plate was coated the day before the analysis, kept at 4°C overnight, and brought to room temperature the morning of the experiment. The coating was done in the tissue culture hood by adding 20 μL of poly-L-Lysine (Sigma P4707) into each well, letting it sit for 10 minutes, aspirating excess coating solution, and letting it dry for 30 minutes. DMEM (Sigma D5030) with 2 mM glutamine, 1 mM pyruvate, 10 mM glucose, and Pen/Strep was used as media during the assay. The pH was adjusted to 7.4 with 1 M NaOH, and the media were sterilized by filtering into an autoclaved container. 400,000 BM or Kit^+^ cells were spun down and washed with 1 mL of the media twice, resuspended in a final volume of 50 μl of media, and loaded in the coated Seahorse plate for analysis. ATP-linked oxygen consumption was calculated as the difference between the average of the 3 measurements before and after Oligomycin injection.

### ATP measurement

Leg bones and spine were collected, cleaned, crushed, and filtered through a 40 μm strainer to collect bone marrow cells which were stained with antibodies against Kit or CD45. Kit^+^ cells were enriched using magnetic selection, stained with other cell surface antibodies, and Lin^-^Kit^+^ HSPCs were sorted as described in the section on Seahorse measurements. Alternatively, unenriched CD45^+^ bone marrow cells were sorted. ∼10,000 cells of each cell type were sorted into 100 μL RPMI-1640 (Gibco, #11835). Cells were kept on ice until measurement using the ATP bioluminescent somatic cell assay kit (Sigma-Aldrich, FLASC), following the manufacturer’s instructions with a few modifications. The procedure was performed in a biosafety cabinet, and no more than 6 samples were processed in parallel to avoid processing time-dependent changes in bioluminescence. 100 μL of ATP Assay Mix working solution was added into each well of a white 96-well plate (Costar, 3917) and allowed to stand for 3 minutes. A different 96-well plate was used to prepare the following mixture for each well: 100 μL ATP-releasing reagent + 50 μL of ultrapure water + 50 μL of cell sample. After mixing, 100 μL of each well was transferred into the white 96-well plate containing ATP assay mix working solution and mixed well by pipetting up and down 3 times. A blank control was prepared by mixing the same reagents except for the cells. The white assay plate was immediately loaded into a microplate reader (Infinite Plex, TECAN) for bioluminescent measurement. The total amount of ATP in moles in each sample was calculated using the equation provided in the assay kit document.

### Metabolite consumption and production analysis

Lin^-^Kit^+^ cells were sorted using the same protocol as for the Seahorse assay (described above). CD45^+^ cells were collected by staining unselected BM cells with CD45-APC for 30 minutes, followed by cell sorting. A total of 500,000 cells were collected per mouse per cell type. Cell suspensions were centrifuged first and washed twice with 1 mL pre-incubation media: pH adjusted (7.4) DMEM (Sigma D5030) with added 2 mM glutamine and 0.2 mM glucose. 50 μL of media was left behind after each wash to minimize cell loss. 50 μL of media was added at the final step for a final volume of 100 μL media + 10 ng/mL SCF, 100 ng/mL TPO, and 1x PSG. The cells were loaded on a 96-well plate, together with media-only blank controls, and incubated in a cell culture incubator for 13 hr. Media were collected by transferring the samples from each well to individual 1.5 mL tubes and centrifuging cells at 4°C, 400 g for 5 minutes. Samples were frozen immediately in liquid nitrogen and stored at -80°C until LC-MS analysis. For MS analysis, 10 μl of the media were added to a 1.5 mL tube, followed by addition of 90 μL of 90% acetonitrile (Optima LC/MS Water W6-4, Acetonitrile A955-4, Fisher Chemical). An internal standard mix, which contained stable isotope-labeled glucose (100 μM), pyruvate (20 μM), lactate (20 μM), glutamine (100 μM), glutamate (10 μM), α-KG (1 μM), succinate (1 μM), malate (1 μM), aspartate (10 μM), and 3-hydroxybutyrate (5 μM) was spiked in (2 μL/sample). The samples were vortexed for 30 sec and incubated on ice 10 minutes, followed by centrifugation at 4°C, 21,000 for 15 minutes. Supernatants were transferred to new Eppendorf tubes, and 30 μL were added to an LC vial with an insert for LC-MS analysis. The injection volume was 15 μL.

### BM-ISF and BM collection for metabolomics

For BM-ISF collection, femur metaphyses were cut and 200 μL DPBS in a 1 mL Luer-Lok syringe and a 25G sterile needle was used to flush one femur, followed by centrifugation at 4°C, 400 g, for 5 min. The supernatant was collected and immediately frozen in liquid nitrogen. To collect BM, pelleted cells were resuspended in 400 μL of ACK buffer for red blood cell lysis for 5 minutes followed by the addition of 1 mL DPBS and centrifugation. The supernatant was aspirated completely, and the cell pellet was immediately frozen in liquid nitrogen. Samples were stored in a -80°C freezer until LC-MS analysis.

### Cell isolation for rare cell metabolomics

The methods used to sort cells for metabolomics and stable isotope tracing analysis were similar to those we reported previously^26,27,37^. Bones were moved to the cold room immediately after cleaning and the remaining procedures were performed in the cold room until sorting. Cells prepared for sorting were in HBSS + 0.2% BSA for metabolomics and DPBS + 0.2% BSA for stable isotope tracing. The sorters were FACS Aria Fusion and FACSymphony S6. 0.5x PBS was used as a sheath fluid and the sorters were operating in 4-way purity mode with a 70 μm nozzle. Cell populations were sorted using the cell surface markers described in other sections. 10,000 cells of each cell type were sorted into 50 μl of 100% acetonitrile and stored in -80^0^C until LC-MS analysis for metabolomics. Blank samples included a ‘test sort’ for 5sec which adds a corresponding volume of sheath fluid passing through the flow cytometer into the blank tube containing the ACN solvent. On the day of analysis, the samples were thawed on ice, vortexed for 1 minute, centrifuged at 21,000 g for 15 minutes at 4^0^C, and transferred to new tubes in a clean PCR hood. 30 μL were transferred to LC-MS vials containing small-volume inserts, and 20 μL were injected into the LC-MS.

### Stable isotope tracing *in vivo*

Mice were fasted for 6-8 hours before the start of infusions and placed under anesthesia using ketamine^37^. Mice were infused with ¹³C₅-L-Glutamine (CLM-1822-H, Cambridge Isotope Laboratories), or ¹⁵N₂-L-Glutamine (NLM-1328, Cambridge Isotope Laboratories), starting with a bolus for 1 minute with 123 μL/min to fill the line with tracer, followed by continuous infusion of 0.0048 mg/g body weight per minute for 4 hours in a volume of 180 μl/hr. Mice were immediately sacrificed, serum was collected from cardiac blood and frozen in liquid nitrogen, and all bones were collected. For rare cell stable isotope tracing, 10,000-15,000 HSC+MPP or HPC1 cells were sorted into 50 μL of 100% acetonitrile, vortexed and stored on dry ice immediately after sorting and transferred to -80^0^C after all samples were sorted. For myeloid progenitors (MyP, Lin^-^Sca-1^-^Kit^+^) 50,000-100,000 cells were sorted into 100μL of DPBS, centrifuged immediately after sorting, most supernatant except for ∼20 μl was removed, and 80 μL 100% was added before storing the samples at -80^0^C. For tracing analysis of total BM, samples were collected as described for the BM collection in the metabolomics section.

### Metabolite extraction for LC-MS analysis

High-purity solvents suitable for UHPLC analysis were used, including Optima LC/MS Water (W6-4, Fisher Chemical) and Acetonitrile (A955-4, Fisher Chemical). For sorted rare cell metabolomics, On the day of the MS analysis, the samples were retrieved and thawed on ice. After vortexing for 1 minute, the samples were placed on ice for 10 minutes, then centrifuged at 21,000 g, 4°C for 15 mins. Supernatants were transferred to LC vials on ice in a clean PCR hood. For LC-MS analysis, 20 μL of the extract was injected. Test-sort samples containing an equal volume of sheath fluid as the samples and solvent-blank samples containing an equal volume of acetonitrile extraction solvent were processed side by side with the experimental samples as negative controls. For total BM cell metabolomics, 150 μL of 80% acetonitrile was added to the cell pellet immediately after removing from the -80°C freezer. Samples were processed with 3 freeze-thaw-vortex cycles (freezing in liquid nitrogen, thawing in ice water) before centrifugation at 21,000 g at 4°C for 15 mins. Supernatants were transferred into a new set of 1.5 mL tubes and centrifuged again and 30 μL were transferred into LC vials containing inserts on ice. 13 μL were injected into the LC-MS. Corresponding sample blank was prepared by adding the same amount of 80% acetonitrile into an empty 1.5 mL tube and processed side by side with the samples. For BM-ISF extraction, 90 μL of 90% acetonitrile was added to 10 μL of BM-ISF sample. After vortexing for 30s, the samples were kept on ice for 10 min before centrifuging at 21,000 g at 4°C for 15 mins. The supernatant was transferred into a new set of 1.5 mL tubes, and 30 μL were transferred to LC-MS vials. A sample blank was prepared by mixing 10 μL DPBS with 90 μL of 90% Acetonitrile and processed the same way as the BM-ISF samples. 15 μL were injected for LC-MS analysis. For serum extraction, 67 μL of 80% acetonitrile was added to 3 μL of serum, vortexed for 30s, left on ice for 10 min, and centrifuged at 21,000 g at 4°C for 15 mins. The supernatant was transferred into a new set of 1.5 mL tubes, centrifuged again, and 30 μL were transferred to LC-MS vials. 10 μL were injected for LC-MS analysis. A sample blank was prepared by mixing 3 μL HPLC-grade water with 67 μL of 80% Acetonitrile and processed the same way as the serum samples. NAD^+^/NADH measurements were performed as previously described^37^, with some modifications. 15,000 cells were sorted into 50 μL ACN:MeOH (v:v) 1:1 with 0.1 M formic acid. Samples were immediately vortexed for 1 minute, neutralized with 32 μL 200 mM NH_4_HCO_3_ (dissolved in ACN:MeOH:H_2_O (v:v:v) 40:40:20) and analyzed immediately on a 6500+ triple quadrupole (Sciex)^37^.

### *Ex vivo* U^13^C-acetate tracing

Bones from euthanized mice were collected, cleaned, and crushed with 10 mL HBSS+0.2% BSA. Cells were filtered through a 40 μm strainer, washed, red blood cells were lysed with ACK lysis buffer, 10 mL HBSS+0.2% BSA were added to stop the lysis, and cells were washed. The BM cell pellet was resuspended in 10 mL DPBS + 0.2% BSA and counted with Vi-Cell XR. 5 million cells were aliquoted per tube and resuspended in 1 ml of DMEM media (Sigma D5030) containing 1 mM U^13^C-sodium acetate, 2 mM glutamine, 1 mM pyruvate, 10 mM glucose, pH 7.4. The samples were incubated in a 37 °C, 5% CO_2_ cell culture incubator for 15 or 60 minutes, centrifuged, and washed with 1 mL of DPBS+0.2% BSA. After aspirating all solution, the cell pellet was immediately frozen in liquid nitrogen and stored at -80°C until MS analysis. For MS analysis, 60 μL of 80% acetonitrile was added, and the metabolite extraction process was the same as the BM cell extraction described above. 13 μL was injected into the LC-MS for analysis.

### LC-MS analysis

To separate metabolites prior to mass spectrometry, a Thermo Scientific (Bremen, Germany) Vanquish Flex liquid chromatography (LC) system was used. LC was performed on a Millipore ZIC-pHILIC column (5 μm, 2.1×150 mm), a binary solvent system of 10 mM ammonium acetate in water, pH 9.8 (solvent A) and acetonitrile (solvent B) with a constant flow rate of 0.25 mL/min or 0.4 mL/min was used. The column was equilibrated with 90% solvent B. The LC gradient was: 0–15 minutes linear ramp from 90% B to 30% B; 15–18 minutes isocratic flow of 30% B; 18–19 minutes linear ramp from 30% B to 90% B; 19–27 column regeneration with isocratic flow of 90% B. Metabolites were analyzed on a Thermo Scientific Orbitrap Exploris 480 mass spectrometer or QExactive HF-X, collecting both positive and negative ion spectra, as previously described^37^. *Cs^Δ/Δ^;Gls^Δ/Δ^* metabolomics analysis was performed on a 6500+ triple quadrupole (Sciex) operating in MRM mode^37^ with the same HILIC column and mobile phase gradient as above. DNA isolation, digestion and LC-MS/MS analysis for 5hmC and 5mC were performed as previously described^64^ using reverse phase LC and a 7500 triple quadrupole (Sciex) in MRM mode.

### Metabolomics data processing

TraceFinder 5.1 (Thermo Scientific) equipped with an in-house spectral library, was used to identify and quantify metabolites. The method compares characterized precursor and product ion spectra to spectra obtained from biological extracts. All metabolites were identified from raw data files with a 5 ppm mass tolerance. Raw values for metabolite MS peak abundance (area under the curve) for the BM, BM-ISF, and nutrient consumption and production experiments were first cleaned by filtering out metabolites for which signal intensity was lower than 3 times the signal intensity in the blank samples. Signal intensities for BM and BM-ISF were normalized to the median signal intensity to account for variability in total sample amount. For *ex vivo* metabolite production data, metabolites were defined as net produced if their average signals were higher than those of the media-only controls. Sorted rare cell metabolomics raw data were first cleaned by filtering out metabolites for which signal intensity was lower than 2 times the signal intensity in the blank sample, and then polar metabolites and non-polar lipids were normalized separately using the corresponding median signal intensity. Metaboanalyst 6.0 was used to process BM metabolomics data and nutrient consumption production data, including missing value imputation (features with missing values in more than 50% of the samples were removed, and for the remaining missing values, a value equal to 0.2 of the minimum value for that feature within the treatment group was used), log transformation, t-tests, false discovery rate correction, and autoscaling for display. Pathway enrichment analysis was done using Metaboanalyst. For stable isotope tracing data, natural abundance was corrected using Accucor^65^.

### Protein Extraction and Western Blot Analysis

For sorted cells, 30,000 CD48^-^Lin^-^Kit^+^Sca-1^+^ or CD48^+^Lin^-^Kit^+^Sca-1^+^ HSPCs were directly sorted into 500 μL of 20% trichloroacetic acid (TCA, Sigma-Aldrich) in Milli-Q H_2_O. 300,000 Lin^-^Kit^+^Sca-1^-^ myeloid progenitors were directly sorted into 50 μL of 20% TCA solution. The final concentration of TCA was adjusted to 10%. Samples were vortexed and centrifuged at 4°C, 21,000 g for 15 min., the precipitate was washed with -20°C cold acetone twice, and dried. Samples were reconstituted in 7 μL of solubilization buffer (2% Triton X-100, 1% DTT, and 9M urea) for HSPCs and 75 μL for myeloid progenitors. 2.5 μL of Laemmli buffer + β-mercaptoethanol was added to HSPC samples, and 25 μL to myeloid progenitor samples. Samples were incubated at 95°C for 10 min or, for membrane proteins at 37°C for 1 hr. Samples were separated on 4-15% Mini-Protean TGX gels (BioRad) and transferred to 0.45 μm or 0.2 μm PVDF membranes (BioRad) by wet transfer using Tris Glycine transfer buffer (BioRad). Western blots were performed using antibodies against CS (Thermo, rabbit polyclonal, PA5-22126), ACLY (Cell Signaling, rabbit polyclonal, D1X6P), ASCT2 (Proteintech, rabbit polyclonal 20350-1-AP), SNAT2 (Proteintech, rabbit polyclonal 25928-1-AP), Vinculin (Cell Signaling, 4650S), β-actin-HRP (Cell Signaling, rabbit monoclonal, 13E5). Signals were detected using the SuperSignal West Pico or SuperSignal West Femto chemiluminescence kits (Thermo). For low numbers of cells, the SuperSignal Western Blot Enhancer kit (Thermo) was used. For total bone marrow, cells were incubated in ACK lysis buffer to lyse red blood cells, followed by protein extraction using RIPA buffer and analysis as above. For gel source data, see Supplementary Figure 1.

### RNA-seq cell isolation and analysis

The bone marrow was collected and Kit^+^ cells were enriched as described above. After cell surface antibody staining and washing with HBSS + 0.1% BSA, CD48^-^Sca-1^+^Kit^+^Lineage^-^ HSCs + MPPs were sorted into 300 μL HBSS+0.1%BSA using yield mode on the sorter. To further purify the population, a 2nd round of sorting was performed, and cells were sorted directly into 300 μL of RLT+1% βME. About 10,000-12,000 were collected per mouse. The samples were vortexed and kept in -80 °C until analysis. RNA extraction and sequencing were performed commercially (Admera Health). RNA was extracted using the RNeasy Micro Kit (Qiagen), quality assessed using TapeStation High Sensitivity RNA ScreenTape Assay (Agilent) and quantified using a Qubit Fluorometer (ThermoFisher). Library construction was performed using SMART-Seq v4 Ultra Low Input RNA Kit (Takara Bio) followed by Nextera XT DNA Library Prep Kit (Illumina) with dual 8-nt indices. Libraries were sequenced on an Illumina Novaseq X Plus 10B generating 35.47 ± 6.19 million 150 bp reverse-stranded paired-end reads. Raw read quality was assessed using FastQC (0.11.8). Raw reads were trimmed using Trim Galore (0.6.4) and were aligned to the Ensembl GRCm38 mouse reference genome using STAR (2.7.9a). Mapped reads were quantified using HTSeq-Count (0.9.1). 16.16 ± 2.73 million exon-mapped reads were normalized, and gene expression levels were measured as fragments per thousand exonic bases per million mapped reads (FPKM) using DESeq2 (1.50.2) with R (4.5.2). To calculate fold-change, the minimum FPKM was set to 0.5. Differential expression tests were performed using DESeq2. Multiple comparisons adjustment was performed using the Benjamini-Hochberg method to control the false discovery rate (FDR). Significantly differentially expressed genes were selected at FDR < 0.05.

### Statistical analysis

To assess the significance of a difference in means between treatments, a t-test or 1-way ANOVA was used when the data did not significantly deviate from normality and did not have significantly unequal variances, a Welch’s t-test or Brown–Forsythe ANOVA when the data did not significantly deviate from normality and had unequal variances, and a Mann–Whitney or Kruskal–Wallis test when the data significantly deviated from normality. To test if data deviated from normality (p < 0.01 for at least one treatment), we used the D’Agostino-Pearson test or, when n < 8, the Shapiro–Wilk test. To test if variance significantly differed among treatments, we used the F-test (for experiments with two treatments) or the Brown–Forsythe test (for more than two treatments). In metabolomics analysis, multiple comparisons correction was performed by controlling the false discovery rate at 5% using the method of Benjamini, Krieger, and Yekutieli. Graphs show *p < 0.05, **p < 0.01, ***p < 0.001 unless noted otherwise. Individual datapoints in all dot plot graphs represent biological replicates (typically, mice). All statistical analysis was performed with measurements from biological replicates. All statistical tests comparing two populations were two-sided. Statistical analyses were performed with Graphpad Prism v10 unless noted otherwise. Graphs were plotted using Prism and heatmaps using Morpheus (Broad).

## Data availability statement

The RNA-seq raw data files are available at NIH SRA PRJNA1482597 and the metabolomics raw data files at Metabolomics Workbench, Study ID ST004971.

## Author contributions

YL and MA designed the project, YL, JR, EOK, XY, JZ, LL, JHJ, and MA performed experiments; YL, JR, and MA analyzed data; ZZ performed bioinformatics analysis; MA directed the project; YL and MA wrote the paper; all authors reviewed the manuscript.

## Acknowledgements

The work was funded by grants from the Cancer Prevention and Research Institute of Texas (CPRIT) (Scholar Award RR180007, RP250280), American Society of Hematology (Faculty Scholar award), Moody Foundation, Welch Foundation (I-2053-20210327), Alex’s Lemonade Stand Foundation (‘A’ Award), the Haggerty Foundation, the Rally Foundation (25IN22), and the National Institutes of Health (R01DK125713, R01HL161387) to MA; an American Society of Hematology Fellow to Faculty Scholar award to YL; and an American Society of Hematology Inclusion Pathway Fellow award to EOK. We thank Hao Zhu, Tripti Sharma and the CRI Mouse Genome Engineering facility for generating the *Cs^fl^* mouse, Tom Mathews and the Children’s Research Institute (CRI) Metabolomics facility (supported by the CPRIT Core Facilities Support Award RP240494 to Ralf DeBerardinis) for mass spectrometry support, the North Texas Clinical Pharmacology Cancer Core (supported by the CPRIT Core Facilities Support Award RP210209 to LL) for 5hmC analysis, Michael Ortiz and the CRI Flow cytometry facility for flow cytometry support, the BioHPC computing cluster at UTSW for computational resources, Dieu Linh Nguyen, Landon Nguyen, and Qing Ding for mouse colony management and technical assistance, Sean Morrison for sharing mice, Courtney Karner for sharing SNAT2 and ASCT2 antibodies, Metafora Biosystems for the gift of GLUT1.RBD, Shawn Burgess and Elvin Wagenblast for discussions, and Prashant Mishra for comments on the manuscript.

---

**Extended Data Figure 1.**
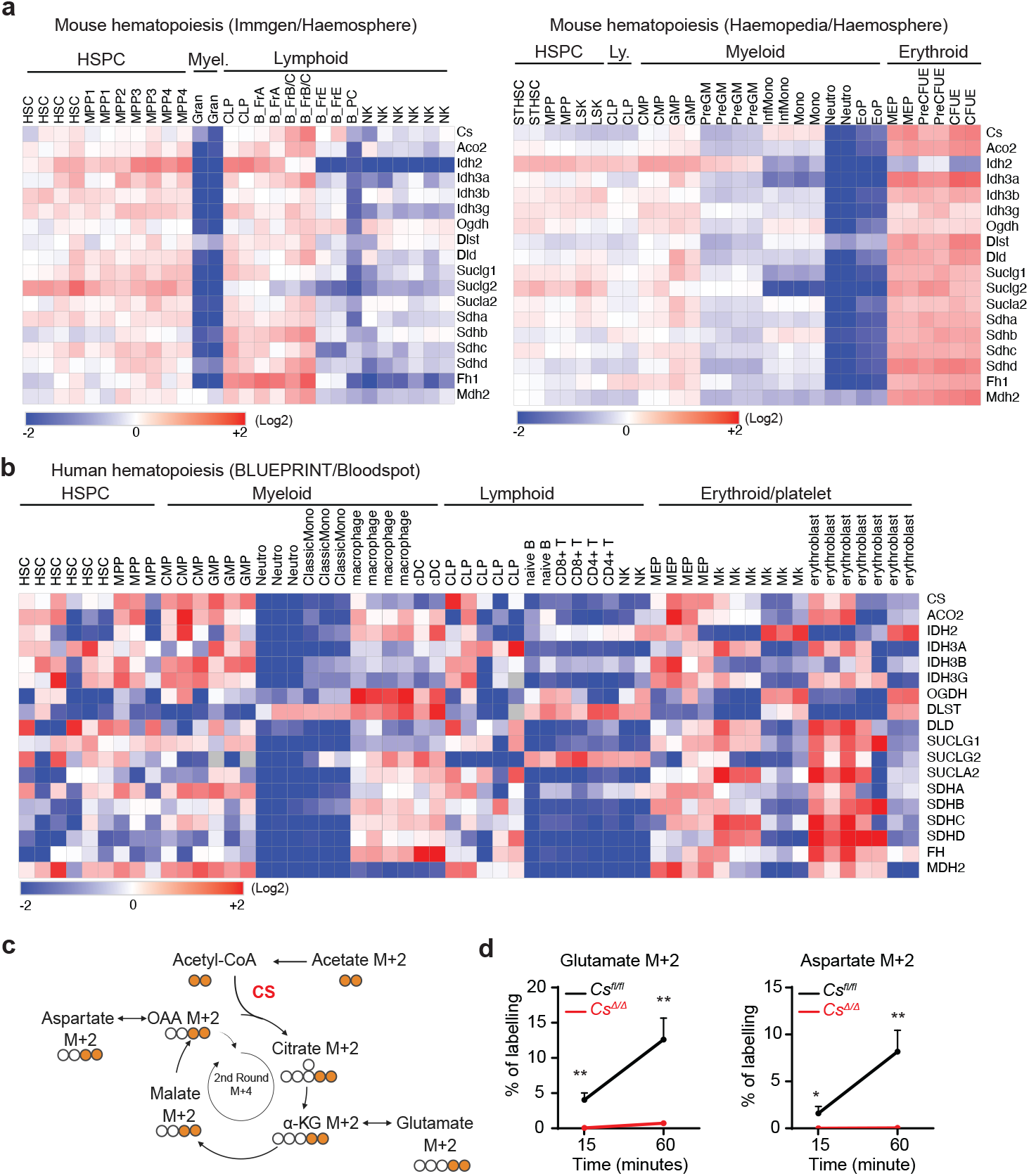
Metabolic and transcriptional changes in the TCA cycle in hematopoietic differentiation. a-b. TCA cycle gene expression levels in mouse and human hematopoietic cell types, shown as relative changes. Data from the Immgen or the Haemopedia RNAseq datasets obtained from the Haemosphere database (b) or the BLUEPRINT RNAseq dataset from the Bloodspot database (c). c-d. Schematic of *ex vivo* tracing of *Mx1Cre;Cs^Δ/Δ^* or littermate control BM cells with U^13^C-acetate to test cycle ablation, and proportion of M+2 labeled glutamate and aspartate. (n = 4-5 mice/genotype) Statistical significance was assessed with a one-way ANOVA of log-transformed data (a), or Welch’s t-test (d).

**Extended Data Figure 2.**
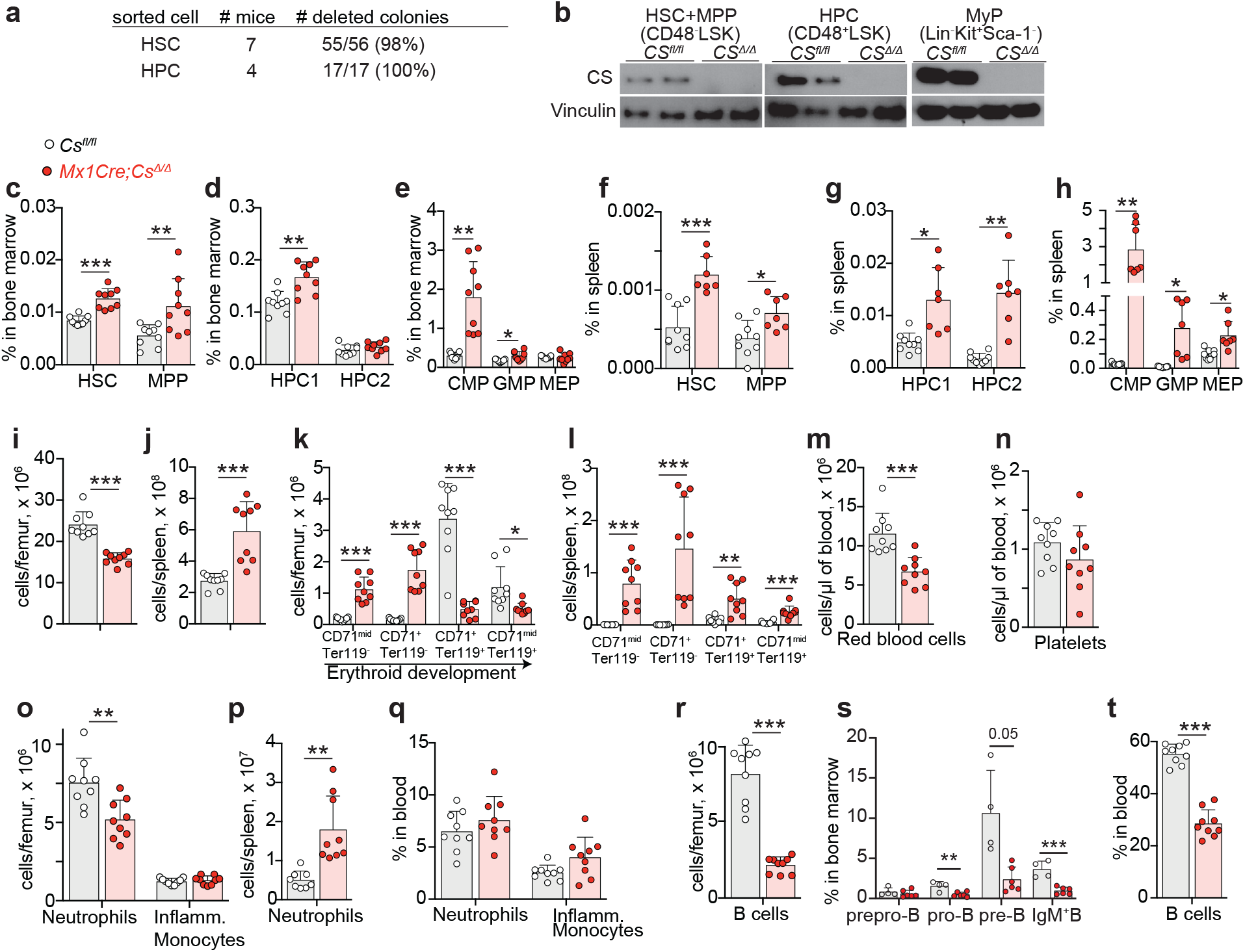
Impact of *Cs* deficiency on hematopoiesis 3 weeks after deletion. a. Genotyping of colonies grown in methylcellulose culture from single HSCs or HPCs sorted from *Mx1Cre;Cs^Δ/Δ^* mice shows that almost all HSCs or HPCs are deleted for *Cs*. b. Western blot results for CS in sorted HSC+MPP, HPC, and Lin^-^Kit^+^Sca-1^-^myeloid progenitors (MyP) from *Mx1Cre;Cs^Δ/Δ^* mice or littermate controls. c-t. Hematopoietic analysis of *Mx1Cre;Cs^Δ/Δ^* mice or littermate *Cs^fl/fl^* controls 3 weeks after poly I:C-mediated deletion. (c-h) frequency of HSCs and progenitors in the BM and spleen; (i-j) number of cells in the bone marrow and spleen; (k-m) number of erythroid lineage cells (n-q) number or frequency of myeloid cells and platelets; (r-s) number or frequency of B cells. Statistical significance was assessed with a t-test (c, d, f, m, o, s-proB, s-IgM^+^B, t), or Welch’s t-test (e, g, h, i, j, k, p, r, s-preB) or Mann-Whitney test (l).

**Extended Data Figure 3.**
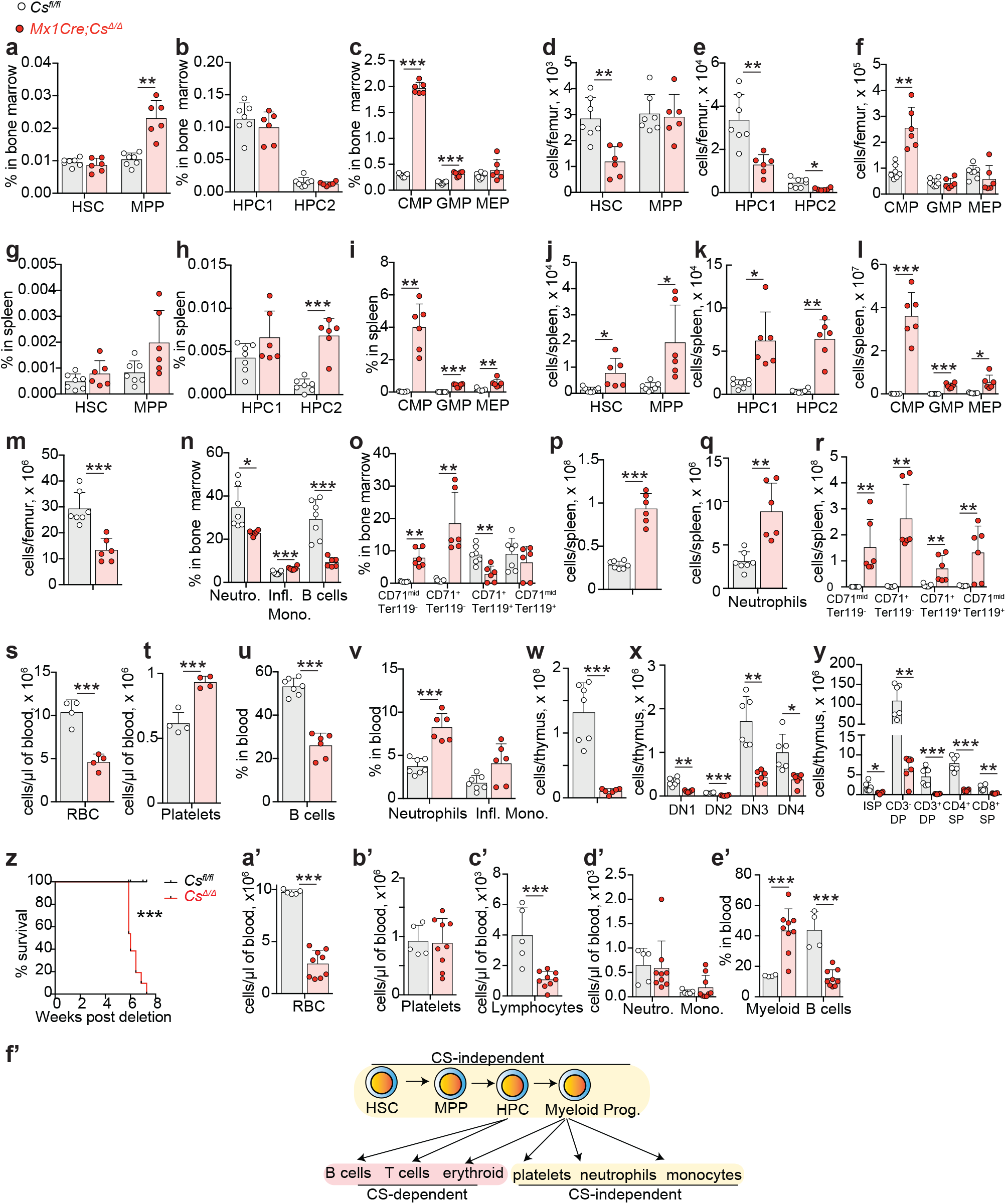
Impact of *Cs* deficiency on hematopoiesis at 5 weeks or at longer periods after deletion. a-υ. Hematopoietic analysis of *Mx1Cre;Cs^Δ/Δ^* mice or littermate *Cs^fl/fl^* controls 5 weeks after poly I:C-mediated deletion. (a-l) frequency and number of HSCs and progenitors in the BM and spleen; (m-r) frequency and number of mature cells in the BM and spleen; (s-v) frequency and number of cells in the blood; (w-y) number of T cell progenitors in the thymus. z. Survival curve of *Mx1Cre;Cs^Δ/Δ^* mice after poly I:C-mediated deletion (n=18-21 mice/genotype). a’-e’. Number or frequency of cells in the blood of moribund *Cs^Δ/Δ^* mice and littermate controls. f’. Schematic summarizing the cell type-specific dependency on CS in hematopoiesis. Statistical significance was assessed with a t-test (d, m, s-v, e’-B cells), or Welch’s t-test (a-c, e-l, n-q, w-y, a’-e’) or Mann-Whitney test (r) or Mantel-Cox test (z).

**Extended Data Figure 4.**
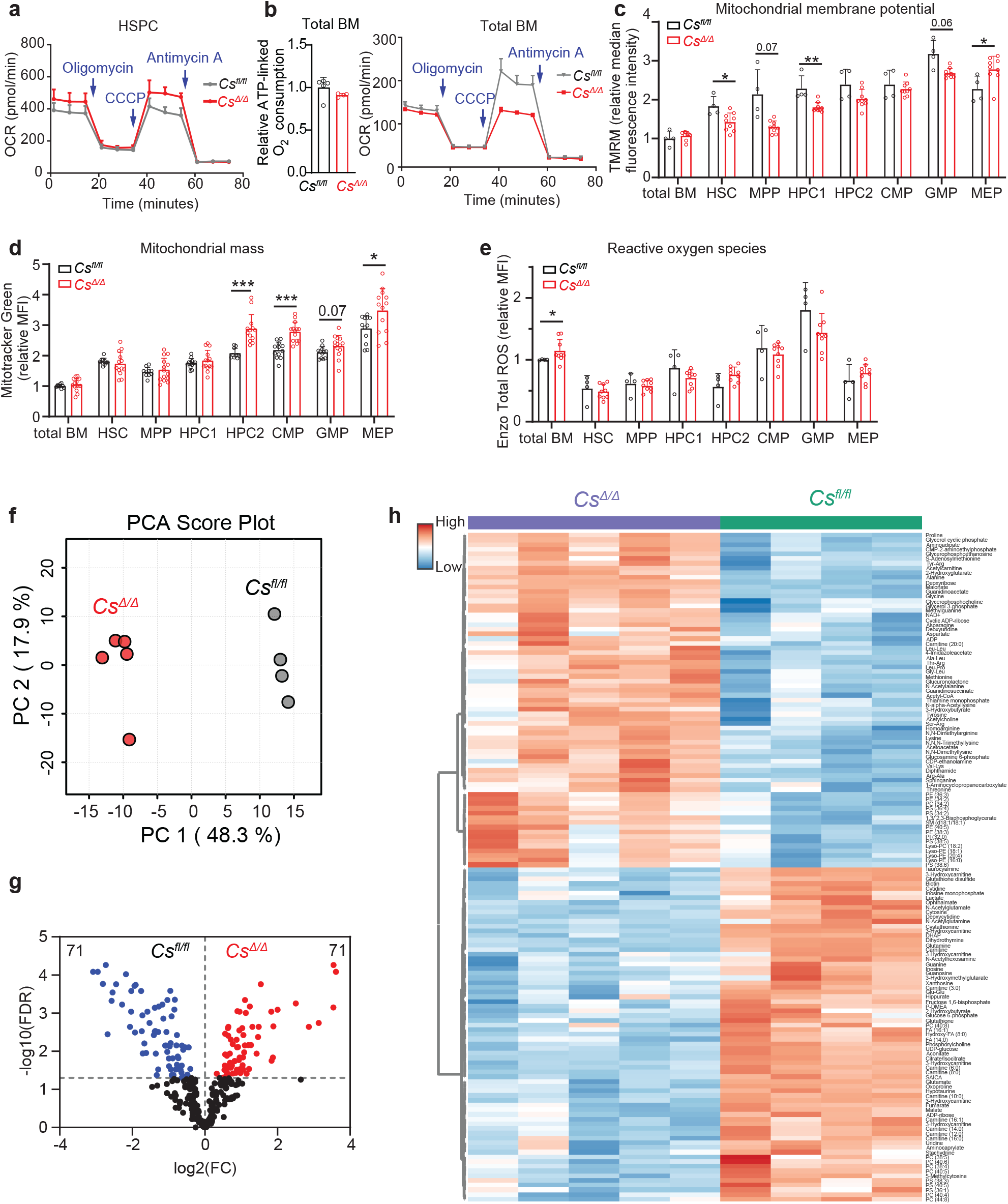
Metabolic analysis of hematopoietic cell types in *Mx1Cre;Cs^Δ/Δ^* mice compared to littermate controls. a. Trace of an oxygen consumption (Seahorse) assay of sorted Lin^-^Kit^+^ HSPCs from *Mx1Cre;Cs^Δ/Δ^* mice and *Cs^fl/fl^* littermate control (the trace shows the average of n = 2 mice/genotype). b. Respiratory O2 consumption of total bone marrow cells and the corresponding trace (n = 4-5 mice/genotype). c-e. Analysis of mitochondrial membrane potential using TMRM (c), mitochondrial mass using Mitotracker Green (d), and ROS using Enzo Total ROS (e) in the indicated cell populations. f-g. PCA analysis (f), volcano plot (g), and heatmap of significantly changed metabolites (h) from total BM metabolomics analysis of *Mx1Cre;Cs^Δ/Δ^* mice and littermate controls. All data represent mean ± s.d. Statistical significance was assessed with a t-test (c-HSC, MEP, d-CMP, GMP), Mann-Whitney test (c-HPC1), Welch’s t-test (c-MPP, GMP, e-total BM), log-transformed t-test (d-HPC2, MEP), or with multiple t-tests on log-transformed data followed by multiple comparisons correction by controlling the FDR at 5% (g-h).

**Extended Data Figure 5.**
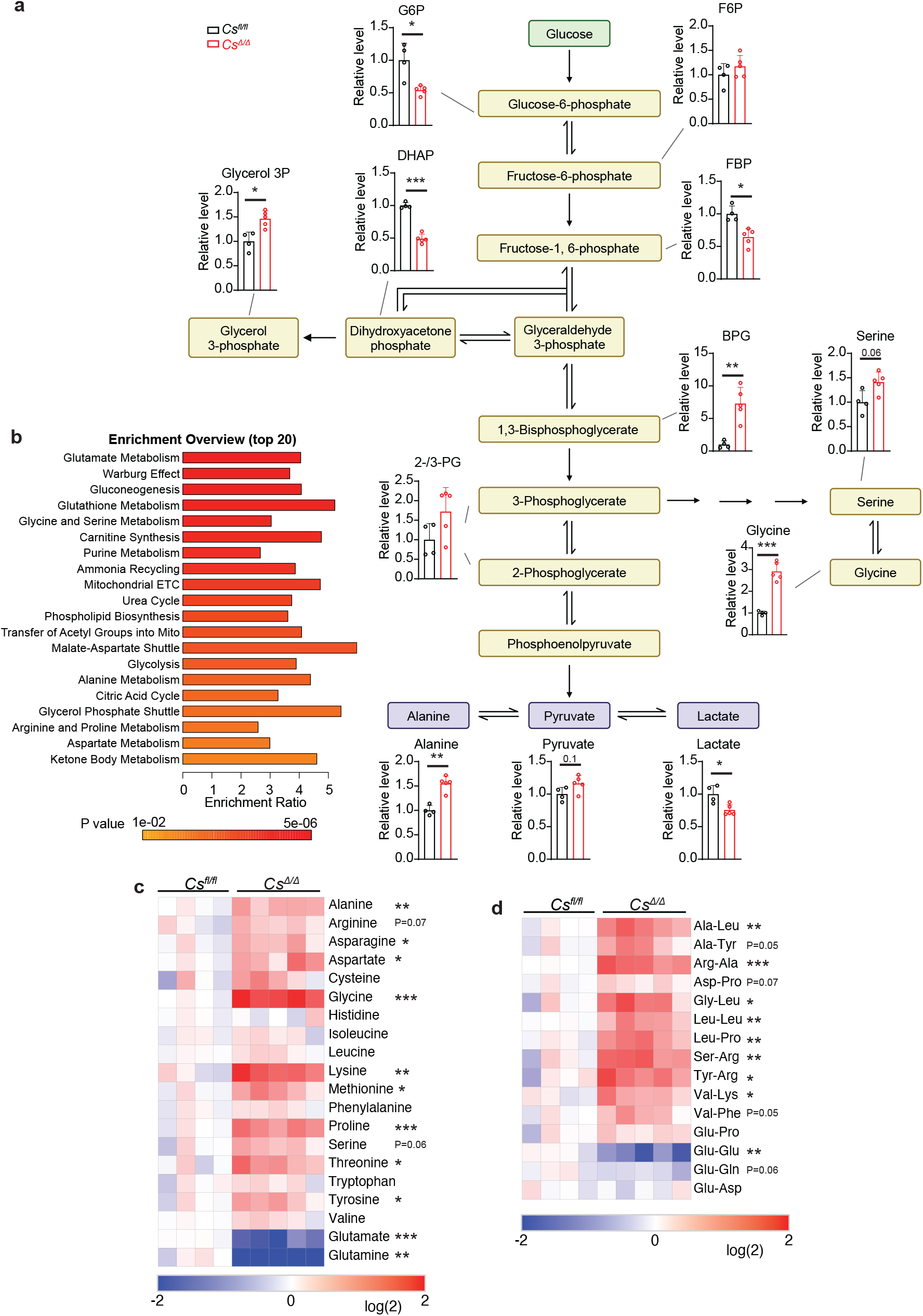
*Cs* deletion induces widespread metabolic changes in glycolysis and amino acid metabolism in bone marrow cells. a. Glycolytic metabolite levels. b. Pathway enrichment analysis of the significantly changed metabolites (FDR < 0.05) showing changes in central carbon and amino acid metabolism. c-d. Effects of *Cs* deletion on amino acids and dipeptide levels. Statistical significance was assessed with a t-test of log-transformed data followed by multiple comparisons correction.

**Extended Data Figure 6.**
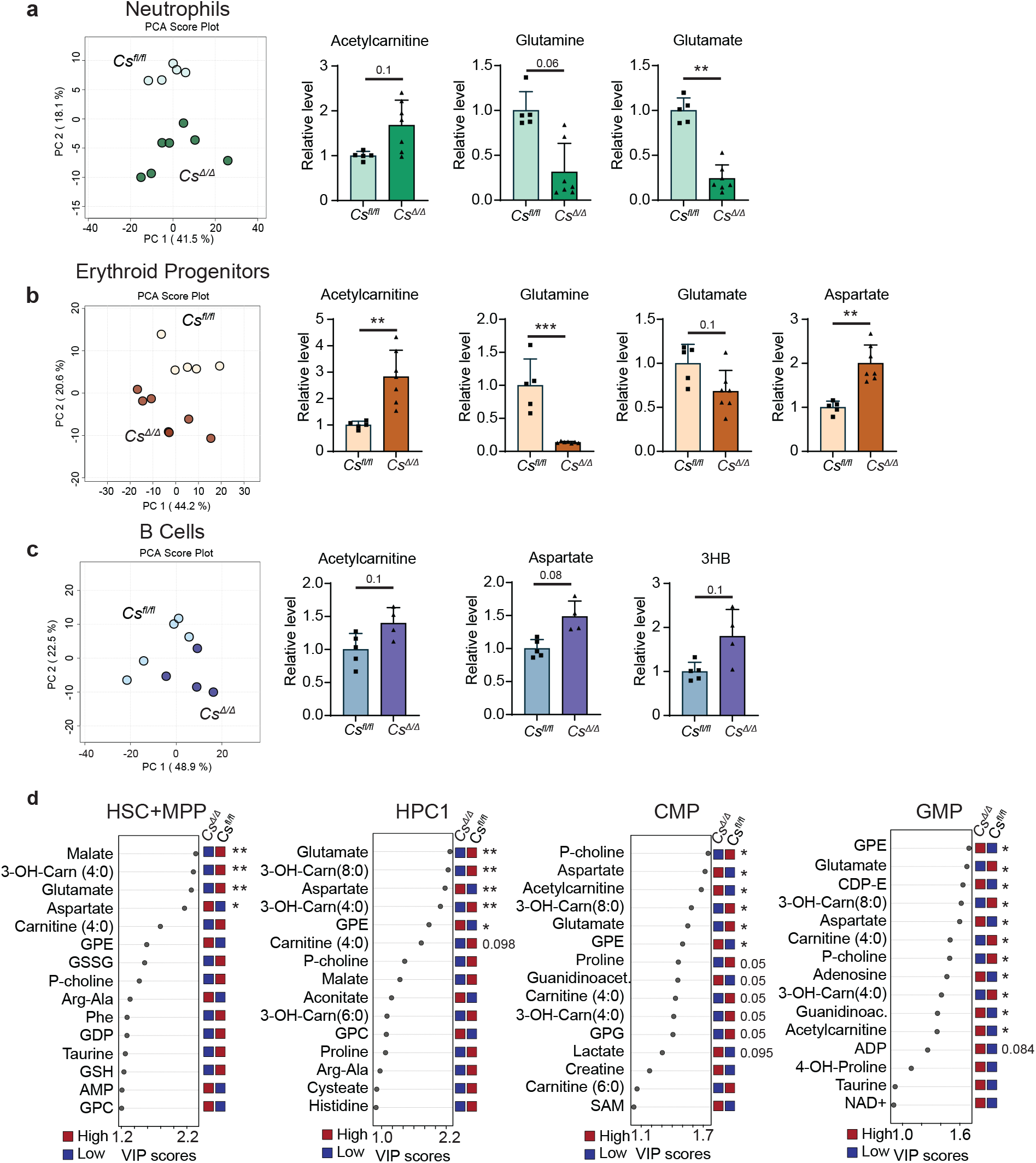
Core metabolic consequences of TCA cycle loss shared across cell types. a-c. PCA analysis of metabolomics data and graphs showing example metabolites that commonly changed in neutrophils (Ly6g^+^), erythroid progenitors (CD71^+^), and B cells (B220^+^) isolated from the bone marrow using magnetic selection. d. PLSDA VIP score plots showing the top 15 differentially abundant polar metabolites in HSC+MPP, HPC1, CMP and GMP cells after *Cs* deletion. Statistical significance was assessed with a t-test of log-transformed data followed by multiple comparisons correction.

**Extended Data Figure 7.**
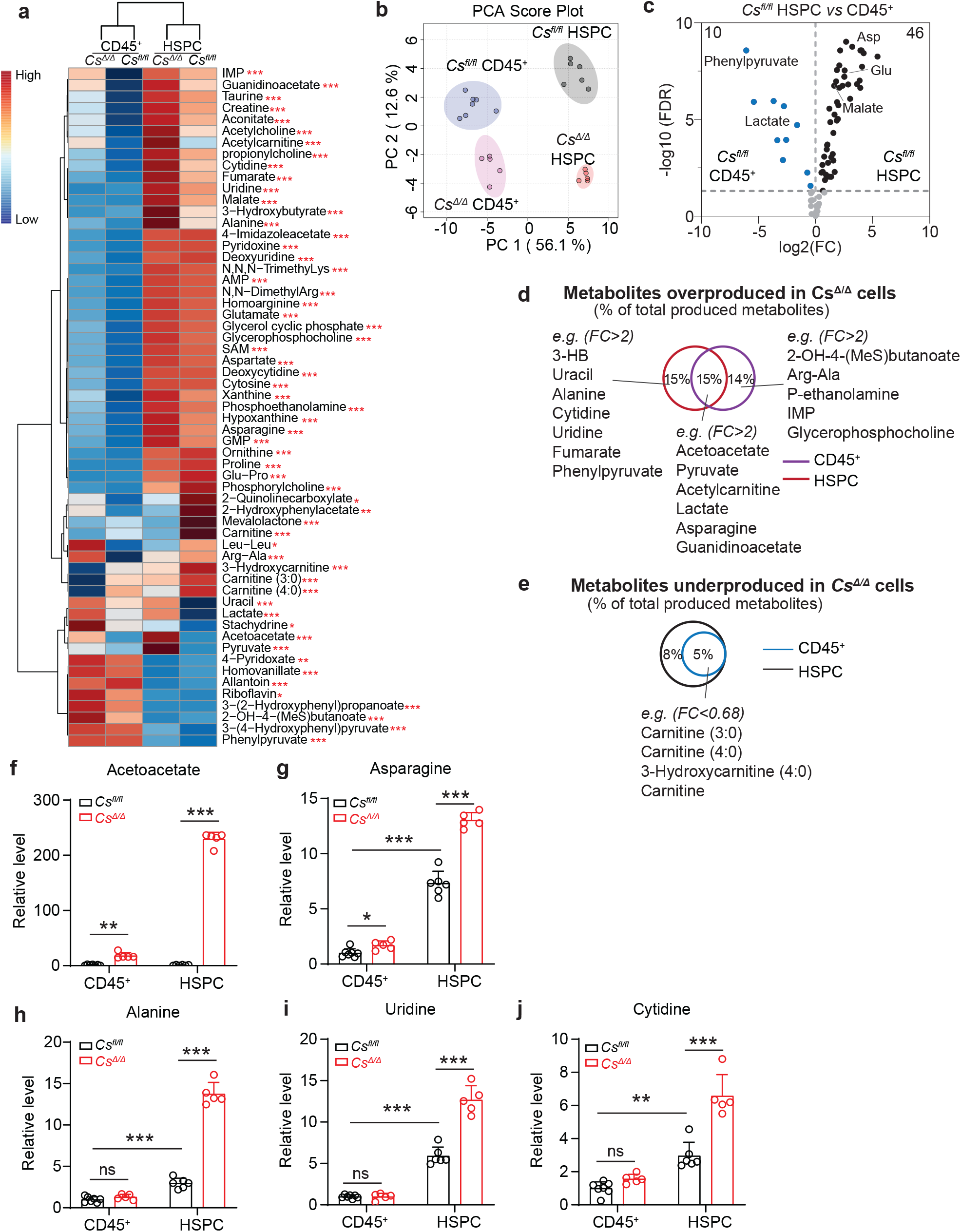
*Cs* deletion changes the metabolite production landscape of total CD45^+^ BM cells and sorted Lin^-^Kit^+^ HSPCs. a. Heatmap of metabolites whose total production significantly changes between cell types or genotypes. b. PCA analysis of the metabolite production landscape in HSPCs and total CD45^+^ BM cells showing that it is driven by both CS activity and cell type. c. Volcano plot showing HSPCs produce more metabolites than CD45+ BM cells. d-e. Metabolites produced at higher (d) or lower (e) rates in *Cs^Δ/Δ^* vs controls HSPCs or CD45^+^ cells. f-j. Metabolites that were significantly overproduced in both total BM and HSPCs (f-g) or only HSPCs (h-j) after Cs deletion. Statistical significance was assessed with one-way ANOVA with log-transformed data followed by multiple comparisons correction.

**Extended Data Figure 8.**
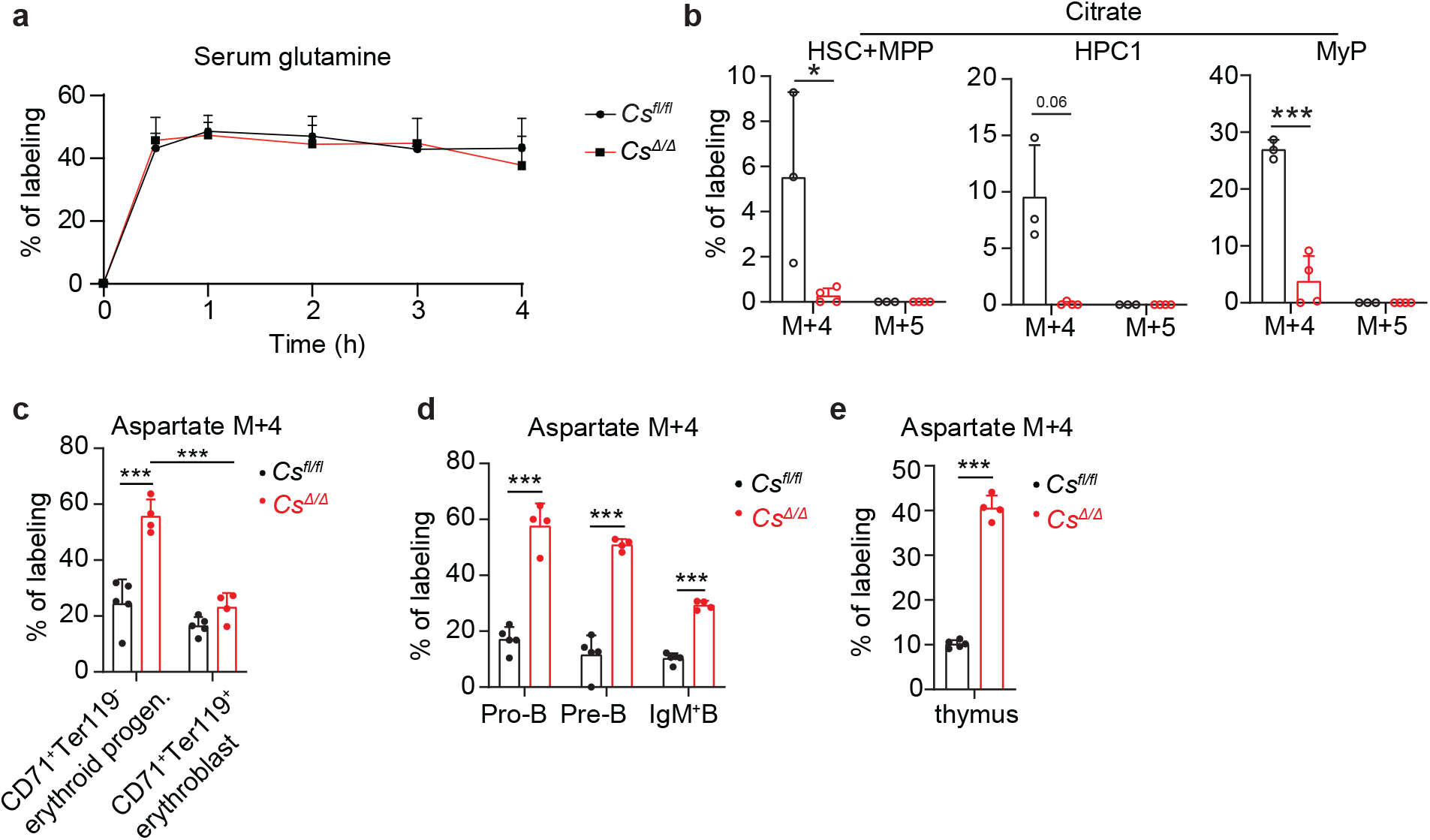
Effects of *Cs* deficiency on U^13^C-Glutamine-derived labeling *in vivo* in different hematopoietic cell types. a. Serum glutamine labeling over time in *Mx1Cre;Cs^Δ/Δ^* mice and littermate controls (n = 3-4 mice/genotype). b. Citrate M+5 labeling. Citrate M+4 labeling (from Fig. 3b) is shown for comparison. c-e. Aspartate M+4 labeling in erythroid lineage cells (c), B cell lineage cells (d), and T cell progenitors in the thymus (e). Statistical significance was assessed with a t-test (c, d), or a Welch’s t-test (e).

**Extended Data Figure 9.**
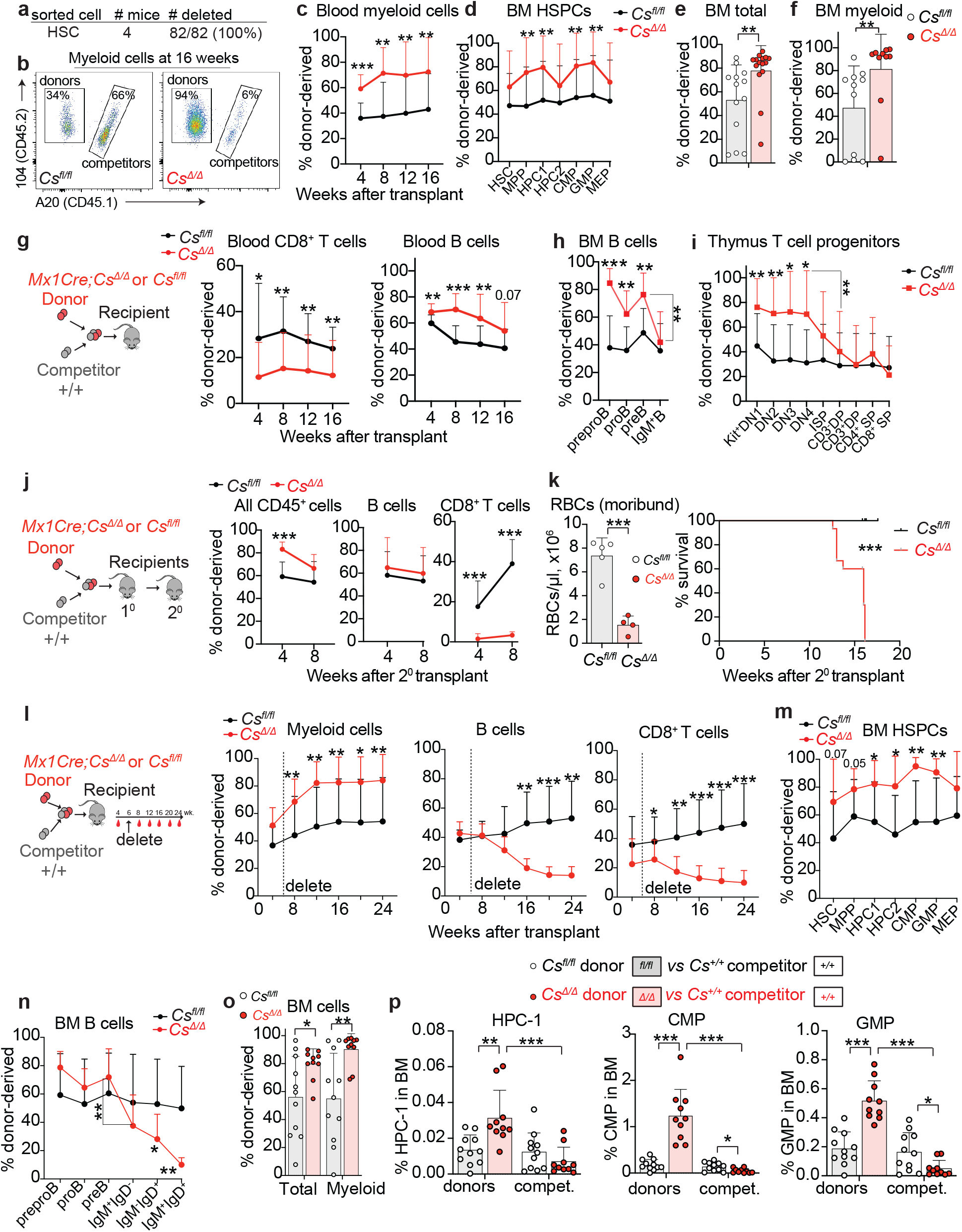
*Cs-*deficiency cell-autonomously increases HSPCs and myeloid reconstitution and impairs lymphoid reconstitution. a. Genotyping of colonies from single donor *Mx1Cre;Cs^Δ/Δ^* HSCs sorted from transplant recipients shows that their deletion efficiency remains high after reconstitution. b. Representative flow plots of myeloid reconstitution in the peripheral blood 16 weeks after competitive transplantation of donor *Mx1Cre;Cs^Δ/Δ^* or *Cs^fl/fl^*bone marrow cells. c-f. Competitive bone marrow transplantation of 500,000 donor *Mx1Cre;Cs^Δ/Δ^* or *Cs^fl/fl^* bone marrow cells with 500,000 wild-type competitor cells into lethally irradiated recipient mice. Mice were from a second *Cs^fl^* line, which was generated from a separate genome-engineered founder mouse. This line was maintained separately from the main *Cs^fl^* line used for the other experiments. Shown are the fraction of donor-derived myeloid cells in the blood (c), of donor-derived HSPCs in the bone marrow (d), and of donor-derived total CD45^+^ or myeloid cells in the bone marrow (e-f) (n=12-15 mice/genotype from 3 independent experiments). g-i. Lymphoid reconstitution after competitive transplantation of 500,000 donor *Mx1Cre;Cs^Δ/Δ^* or *Cs^fl/fl^* bone marrow cells with 500,000 wild-type competitor cells as shown in Fig. 4a. Donor-derived reconstitution of CD8^+^ T cells and B cells in the blood (g), B cell progenitors in the bone marrow (h), and T cell progenitors in the thymus (i) (n=13-15 mice/genotype for blood analysis and 10-13 mice/genotype for bone marrow or thymus analysis from 3 independent experiments). j-k. Secondary transplantation of 5 million bone marrow cells from primary competitive transplant recipients. (j) total, B, and T cell reconstitution in the blood (n=15 mice/genotype). (k) Anemia and mortality in secondary transplant recipient mice (n=15 mice/genotype). l-p. Competitive bone marrow transplantation of 500,000 donor *Mx1Cre;Cs^ΔlΔ^* or *Cs^fl/fl^* bone marrow cells with 500,000 wild-type competitor cells into lethally irradiated recipient mice followed by poly I:C 6 weeks post-transplant. Shown are the fraction of donor-derived myeloid, B and T cells in the blood (l), HSPC populations in the bone marrow (m), B cell progenitors and mature B cells in the bone marrow (n), total and myeloid cells in the bone marrow (o), and the percentage of donor or competitor progenitor cells out of total bone marrow cells (p). (n=15 mice/genotype for blood analysis and 11 mice/genotype for bone marrow analysis from 3 independent experiments). Statistical significance was assessed with a Mann-Whitney test (c-f, g-T cells, h-j, l-T cells, l-myeloid, m-HPC1, HPC2), t-test (g-B cells, m-HSC, MPP, o-*Cs^Δ/Δ^* vs *CS^fl/fl^* comparison), Welch’s t-test (k-RBCs, l-B cells, m-CMP, GMP, n, o), Mantel-Cox test (k-survival curve), Wilcoxon matched-pairs signed rank test (h,n-preB vs IgM^+^ B cell comparison, i-DN4 vs CD3^-^DP cell comparison), and paired t-test (p-donor vs competitor comparisons).

**Extended Data Figure 10.**
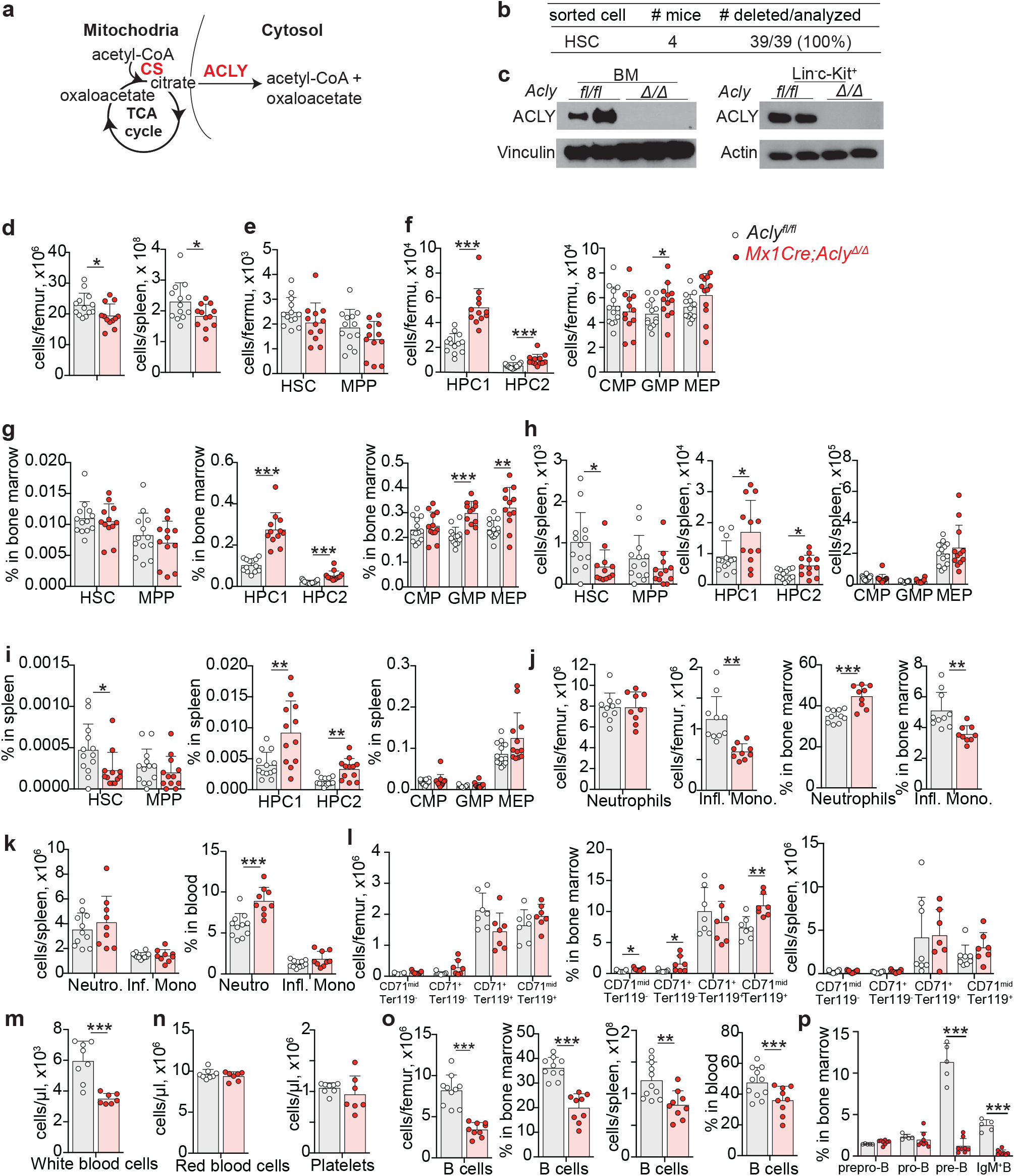
*Acly* deletion does not phenocopy *Cs* deletion at 3 weeks post-deletion. a. Illustration of the CS-ACLY pathway for cytosolic citrate use. a. Genotyping to assess *Acly* deletion of colonies from single HSCs sorted from *Mx1Cre;Acly^fl/fl^* mice ∼ 1 year after poly I:C administration. c. Western blot showing ACLY depletion in *Mx1Cre;Acly^Δ/Δ^* BM cells and sorted Lin^-^Kit^+^ HSPCs. d-p. Hematopoietic analysis of *Mx1Cre;Acly^Δ/Δ^* mice or littermate *Acly^fl/fl^* controls 3 weeks after poly I:C administration. Shown are: (d) bone marrow and spleen cellularity; (e-i) frequency and number of HSCs and progenitors in the BM and spleen; (j-p) frequency or number of: (j-k) myeloid cells, (l) erythroid cells, (m-n) leukocytes, erythrocytes or platelets, and (o-p) B lineage cells in the bone marrow, spleen, or blood. Statistical significance was assessed with a t-test (d, f-GMP, g-GMP, MEP, o, j, k, p-IgM^+^B), Mann-Whitney test (f-HPC1, HPC2, g-HPC1, HPC2, h-HSC, i-HSC, l), and Welch’s t-test (h-HPC1, HPC2, i-HPC1, HPC2, m, p-pre-B).

**Extended Data Figure 11.**
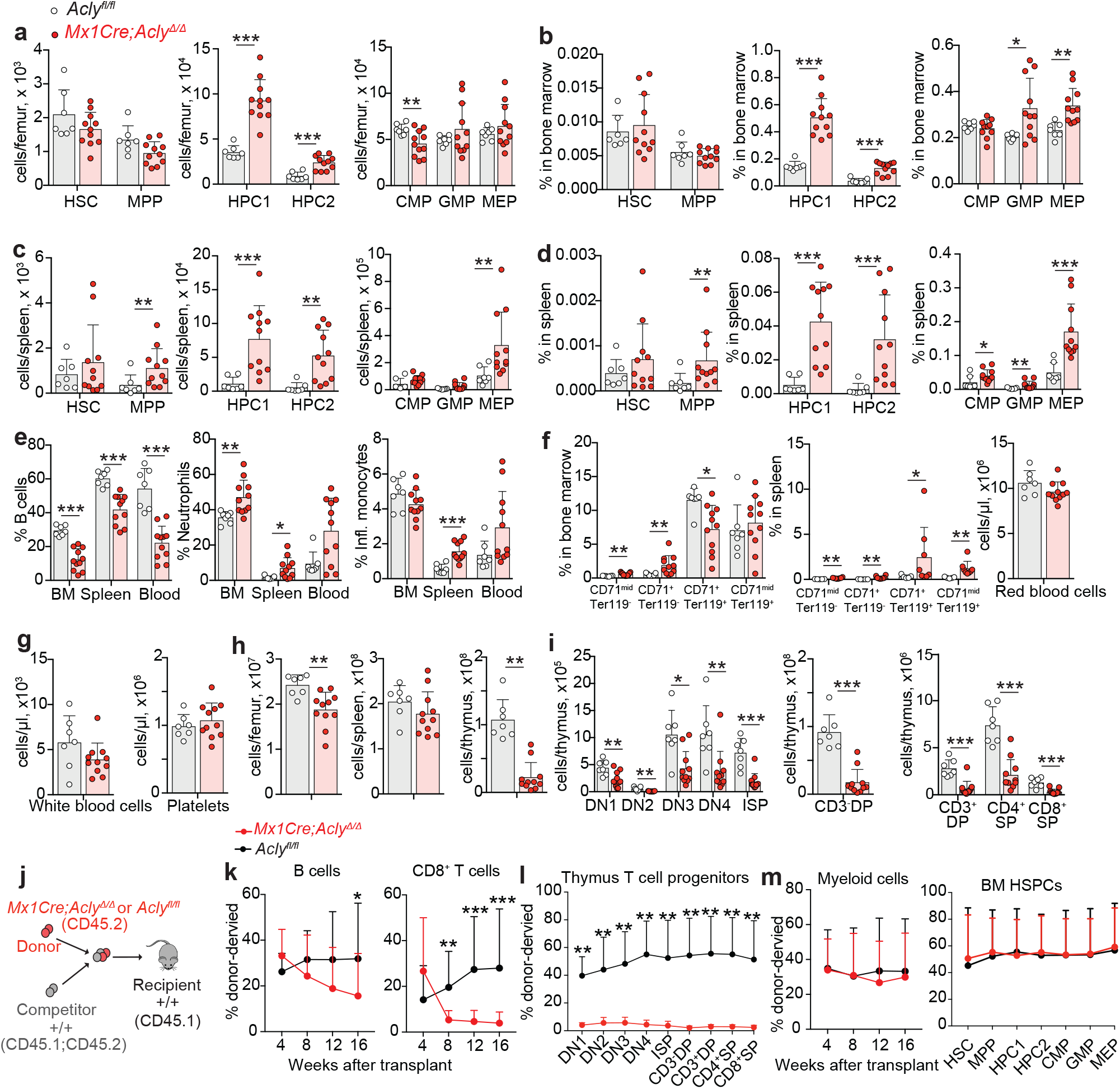
Effects of *Acly* deletion on hematopoiesis at 8 weeks post-deletion and on competitive transplantation. a-h. Hematopoietic analysis of *Mx1Cre;Acly^Δ/Δ^*mice or littermate *Acly^fl/fl^* controls 8 weeks after poly I:C-mediated deletion. Shown are: (a-d) frequency and number of HSCs and progenitors in the BM and spleen; (e-f) frequency or number in the BM, spleen, or blood of B or myeloid cells (e) and erythroid cells (f); (g) white blood cell and platelet counts; (h) BM, spleen, and thymus cellularity; (i) numbers of T cell progenitors in the thymus. j-m. Competitive bone marrow transplantation of 500,000 donor *Mx1Cre;Acly^Δ/Δ^*or *Acly^fl/fl^* bone marrow cells with 500,000 wild-type competitor cells into lethally irradiated recipient mice 3 weeks after poly I:C-mediated deletion. Shown are the fraction of donor-derived B and T cells in the blood (k), T cell progenitors in the thymus (l), myeloid cells in the blood and HSPCs in the bone marrow (m) (n=13-15 mice/genotype for blood analysis, 7-9 for bone marrow analysis and 6-8 mice for thymus analysis from 3 independent experiments). Statistical significance was assessed with a Welch’s t-test (a, b, e-Neutrophils, f-BM, i, l), Mann-Whitney test (c, d, f-spleen, k), and t-test (e-B cells, monocytes, h).

**Extended Data Figure 12.**
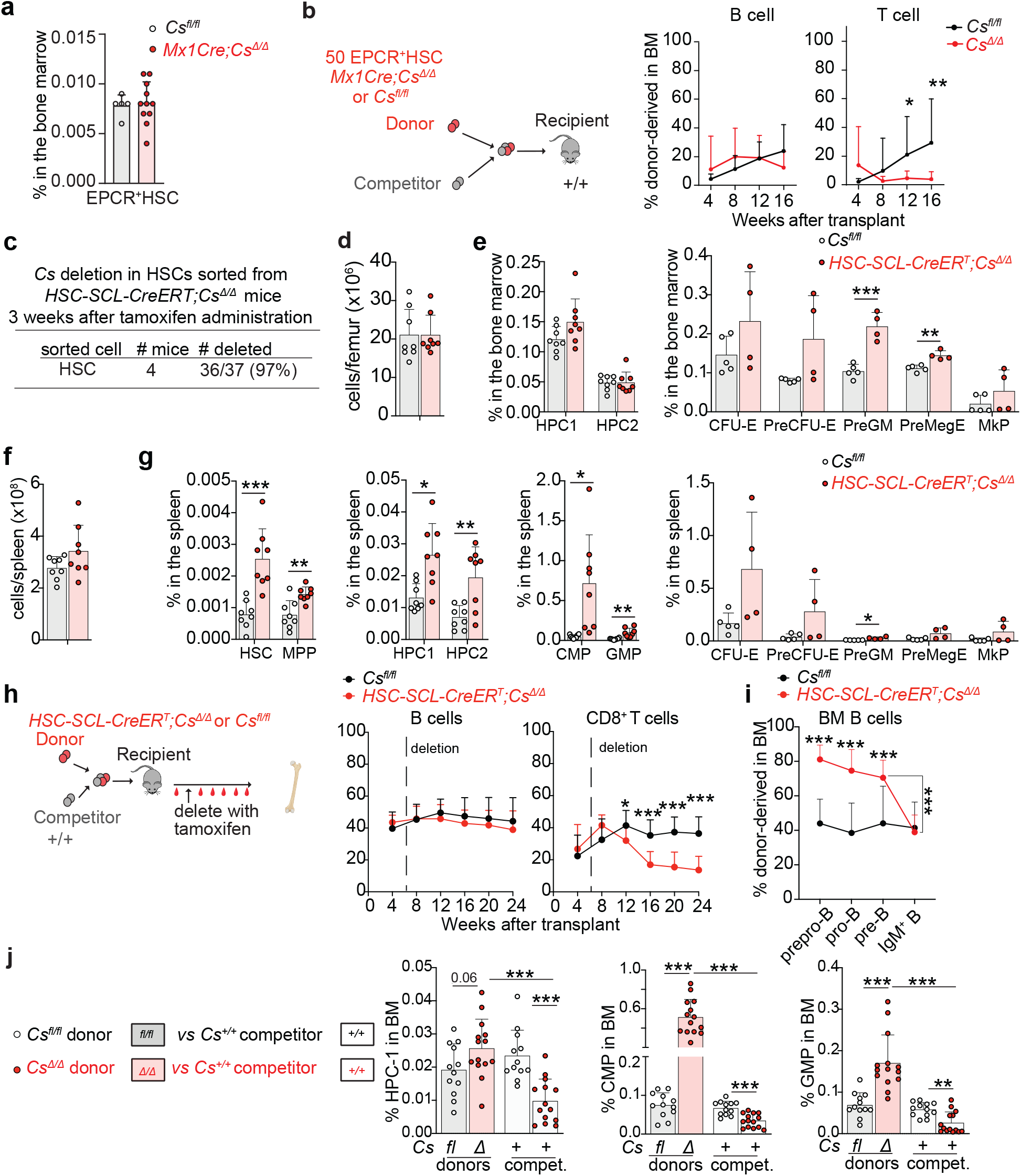
Purified EPCR^+^HSC transplants or *HSC-Scl-CreER^T^*-driven deletion experiments show that *Cs* deletion cell-autonomously promotes HSC function and impairs lymphoid differentiation. a. Frequency of EPCR^+^HSCs in the BM from *Mx1Cre;Cs^Δ/Δ^* mice or littermate *Cs^fl/fl^* controls 3 weeks after poly I:C-mediated deletion. b. Competitive bone marrow transplantation of 50 purified donor EPCR^+^HSCs from *Mx1Cre;Cs^Δ/Δ^* or *Cs^fl/fl^* mice with 1,000,000 wild-type total bone marrow competitor cells into lethally irradiated recipient mice. Shown are the fractions of donor-derived B and T cells in the blood (n=9-12 mice/genotype from 3 experiments). c. Genotyping to assess *Cs* deletion of colonies derived from single HSCs sorted from *HSC-Scl-CreER^T^;Cs^fl/fl^* mice 3 weeks after tamoxifen administration. d-g. Hematopoietic analysis of *HSC-SCL-CreER^T^;Cs^Δ/Δ^* mice or littermate *Cs^fl/fl^* controls 3 weeks after tamoxifen-mediated deletion. (d) BM cellularity; (e) frequency of progenitors in the BM; (f) cell number in the spleen; (g) frequency of HSCs and progenitors in the spleen. h-j. Competitive bone marrow transplantation of 2,000,000 donor *HSC-SCL-CreER^T^;Cs^fl/fl^*or *Cs^fl/fl^* bone marrow cells with 2,000,000 wild-type competitor cells into lethally irradiated recipient mice followed by tamoxifen administration 6 weeks post-transplant. Shown are the fraction of donor-derived B and T cells in the blood (h), of donor-derived B cell progenitor cells in the bone marrow (i), and the percentage of donor or competitor progenitor cells out of total bone marrow cells (j) (n=13-15 mice/genotype for blood analysis and 12-14 mice/genotype for bone marrow analysis from 3 independent experiments). Statistical significance was assessed with a t-test (e, g-HSC, MPP, HPC1, HPC2, h, i, j-HPC1 donor and HPC1, CMP, GMP competitor *Cs^Δ/Δ^* vs *Cs^fl/fl^* comparisons), Mann-Whitney test (b), Welch’s t-test (g-CMP, GMP, preGM, j-CMP, GMP donor *Cs^Δ/Δ^* vs *Cs^fl/fl^* comparisons), and paired t-test (i-preB vs IgM^+^ B, j-donor vs competitor comparisons).

**Extended Data Figure 13.**
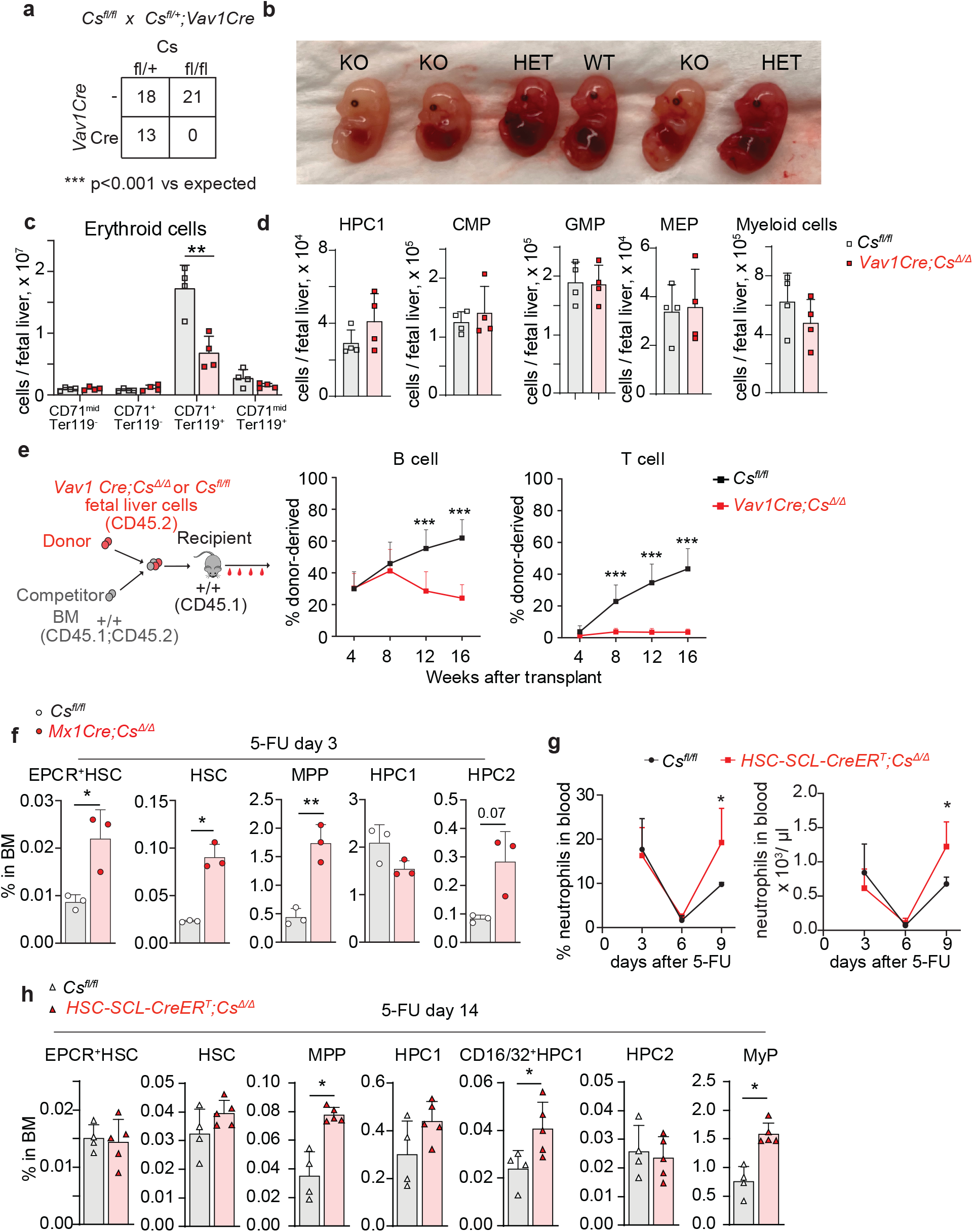
*Cs* deletion promotes fetal liver HSC function and adult hematopoietic regeneration after 5-FU. a. Genotypes of P0 mice born from the indicated *Cs^fl/fl^ x Cs^fl/+^;Vav1Cre* cross. b. Photos of E14.5 embryos from the same cross as (a). WT= *Cs^fl/fl^* or *Cs^fl/+^,* HET= *Cs^fl/+^;Vav1Cre*, KO=*Cs^fl/fl^;Vav1Cre*. c-d. Hematopoietic analysis of the fetal liver of E14.5 *Vav1Cre;Cs^Δ/Δ^* embryos or littermate controls (*Cs^fl/fl^* or *Cs^fl/+^*). The number of erythroid cells (c), HSPCs, and myeloid cells (d) is shown. e. Competitive bone marrow transplantation of 500,000 donor *Vav1Cre;Cs^Δ/Δ^* or littermate control fetal liver cells with 2,000,000 wild-type competitor total BM cells into lethally irradiated recipient mice. Shown are the fractions of donor-derived B and T cells in the blood (n=14 mice/genotype from 3 independent experiments). f. Frequency of HSPCs in *Mx1Cre;Cs^Δ/Δ^* mice or controls 3 days after 5-FU. g. Recovery of blood neutrophils after 5-FU in *HSC-SCL-CreER^T^;Cs^Δ/Δ^* mice or controls (n=4-5 mice/genotype). h. Frequency of HSPCs in *HSC-SCL-CreER^T^;Cs^Δ/Δ^* mice or controls 14 days after 5-FU. All data represent mean ± s.d. Statistical significance was assessed with a chi-square test (a), t-test (c-CD71^+^Ter119^+^, e-B cells, f-EPCR^+^HSC and MPP, g, h-CD16/32^+^HPC1), Mann-Whitney test (h-MyP), or Welch’s t-test (e-T cells, f-HPC2, h-MPP).

**Extended Data Figure 14.**
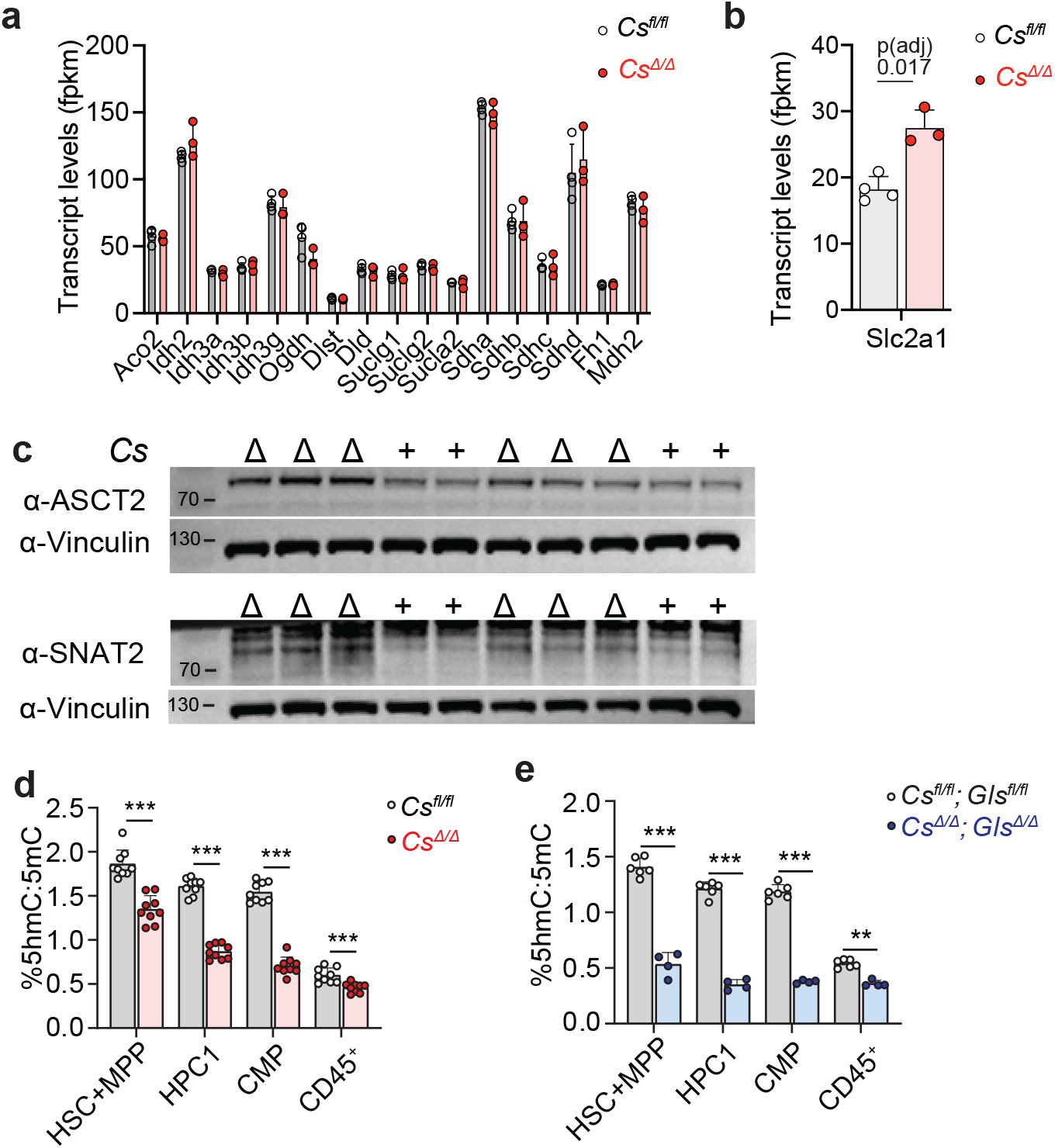
Effects of *Cs* deletion on gene expression, nutrient transporter levels and 5hmC levels. a-b. Expression levels of TCA cycle genes and *Slc2a1* in RNA-seq analysis of *Mx1Cre;Cs^fl/fl^* or control HSC+MPP. c. Western blot of glutamine transporter levels in sorted HSPCs. The ∼75 KDa band is quantified in Fig. 6. d-e. 5hmC/5mC ratio in hydrolyzed DNA of sorted HSPC populations from the indicated genotypes. All data represent mean ± s.d. Statistical significance was assessed with a t-test (d, e) or in RNA-seq with DESeq2 followed by the Benjamini-Hochberg method to control the false discovery rate.

## Notes

### Competing Interest Statement

The authors have declared no competing interest.

## References.

1 Krebs, H. A. & Johnson, W. A. The role of citric acid in intermediate metabolism in animal tissues. Enzymologia 4 (1937).

2 Krebs, H. A. Rate control of the tricarboxylic acid cycle. Adv Enzyme Regul 8, 335–353 (1970). 10.1016/0065-2571(70)90028-2

3 Krebs, H. A. The Citric Acid Cycle. Nobel Prize Lecture (1953).

4 Krebs, H. A. Reminiscences and Reflections. (Oxford University Press, 1982).

5 Haslam, R. J. & Krebs, H. A. The Metabolism of Glutamate in Homogenates and Slices of Brain Cortex. Biochem J 88, 566–578 (1963). 10.1042/bj0880566

6 Chandel, N. S. Mitochondria. Cold Spring Harb Perspect Biol 13 (2021). 10.1101/cshperspect.a040543

7 Inigo, M., Deja, S. & Burgess, S. C. Ins and Outs of the TCA Cycle: The Central Role of Anaplerosis. Annu Rev Nutr 41, 19–47 (2021). 10.1146/annurev-nutr-120420-025558

8 Agathocleous, M. et al. Metabolic differentiation in the embryonic retina. Nat Cell Biol 14, 859–864 (2012). 10.1038/ncb2531

9 Agathocleous, M. & Harris, W. A. Metabolism in physiological cell proliferation and differentiation. Trends Cell Biol 23, 484–492 (2013). 10.1016/j.tcb.2013.05.004

10 Meacham, C. E., DeVilbiss, A. W. & Morrison, S. J. Metabolic regulation of somatic stem cells in vivo. Nat Rev Mol Cell Biol 23, 428–443 (2022). 10.1038/s41580-022-00462-1

11 Warburg, O. On the origin of cancer cells. Science 123, 309–314 (1956).

12 Bartman, C. R. et al. Slow TCA flux and ATP production in primary solid tumours but not metastases. Nature 614, 349–357 (2023). 10.1038/s41586-022-05661-6

13 Sender, R., Fuchs, S. & Milo, R. Revised Estimates for the Number of Human and Bacteria Cells in the Body. PLoS Biol 14, e1002533 (2016). 10.1371/journal.pbio.1002533

14 Sender, R. & Milo, R. The distribution of cellular turnover in the human body. Nat Med 27, 45–48 (2021). 10.1038/s41591-020-01182-9

15 Simsek, T. et al. The distinct metabolic profile of hematopoietic stem cells reflects their location in a hypoxic niche. Cell Stem Cell 7, 380–390 (2010). 10.1016/j.stem.2010.07.011

16 Vannini, N. et al. Specification of haematopoietic stem cell fate via modulation of mitochondrial activity. Nat Commun 7, 13125 (2016). 10.1038/ncomms13125

17 Liang, R. et al. Restraining Lysosomal Activity Preserves Hematopoietic Stem Cell Quiescence and Potency. Cell Stem Cell 26, 359–376 e357 (2020). 10.1016/j.stem.2020.01.013

18 Halvarsson, C., Eliasson, P. & Jonsson, J. I. Pyruvate dehydrogenase kinase 1 is essential for transplantable mouse bone marrow hematopoietic stem cell and progenitor function. PLoS One 12, e0171714 (2017). 10.1371/journal.pone.0171714

19 Takubo, K. et al. Regulation of glycolysis by Pdk functions as a metabolic checkpoint for cell cycle quiescence in hematopoietic stem cells. Cell Stem Cell 12, 49–61 (2013). 10.1016/j.stem.2012.10.011

20 Vannini, N. et al. The NAD-Booster Nicotinamide Riboside Potently Stimulates Hematopoiesis through Increased Mitochondrial Clearance. Cell Stem Cell 24, 405–418 e407 (2019). 10.1016/j.stem.2019.02.012

21 Qiu, J. et al. Using mitochondrial activity to select for potent human hematopoietic stem cells. Blood Adv 5, 1605–1616 (2021). 10.1182/bloodadvances.2020003658

22 Mantel, C. R. et al. Enhancing Hematopoietic Stem Cell Transplantation Efficacy by Mitigating Oxygen Shock. Cell 161, 1553–1565 (2015). 10.1016/j.cell.2015.04.054

23 Bejarano-Garcia, J. A. et al. Sensitivity of hematopoietic stem cells to mitochondrial dysfunction by SdhD gene deletion. Cell Death Dis 7, e2516 (2016). 10.1038/cddis.2016.411

24 Guitart, A. V. et al. Fumarate hydratase is a critical metabolic regulator of hematopoietic stem cell functions. J Exp Med 214, 719–735 (2017). 10.1084/jem.20161087

25 Anso, E. et al. The mitochondrial respiratory chain is essential for haematopoietic stem cell function. Nat Cell Biol 19, 614–625 (2017). 10.1038/ncb3529

26 Agathocleous, M. et al. Ascorbate regulates haematopoietic stem cell function and leukaemogenesis. Nature 549, 476–481 (2017). 10.1038/nature23876

27 DeVilbiss, A. W. et al. Metabolomic profiling of rare cell populations isolated by flow cytometry from tissues. Elife 10 (2021). 10.7554/eLife.61980

28 Passegue, E., Wagers, A. J., Giuriato, S., Anderson, W. C. & Weissman, I. L. Global analysis of proliferation and cell cycle gene expression in the regulation of hematopoietic stem and progenitor cell fates. J Exp Med 202, 1599–1611 (2005). 10.1084/jem.20050967

29 Eastman, A. E. et al. Resolving Cell Cycle Speed in One Snapshot with a Live-Cell Fluorescent Reporter. Cell Rep 31, 107804 (2020). 10.1016/j.celrep.2020.107804

30 Cappel, D. A. et al. Pyruvate-Carboxylase-Mediated Anaplerosis Promotes Antioxidant Capacity by Sustaining TCA Cycle and Redox Metabolism in Liver. Cell Metab 29, 1291–1305 e1298 (2019). 10.1016/j.cmet.2019.03.014

31 Li, X. et al. Circulating metabolite homeostasis achieved through mass action. Nat Metab 4, 141–152 (2022). 10.1038/s42255-021-00517-1

32 Watanuki, S. et al. Context-dependent modification of PFKFB3 in hematopoietic stem cells promotes anaerobic glycolysis and ensures stress hematopoiesis. Elife 12 (2024). 10.7554/eLife.87674

33 Garcia-Bermudez, J. et al. Aspartate is a limiting metabolite for cancer cell proliferation under hypoxia and in tumours. Nat Cell Biol 20, 775–781 (2018). 10.1038/s41556-018-0118-z

34 Qi, L. et al. Aspartate availability limits hematopoietic stem cell function during hematopoietic regeneration. Cell Stem Cell 28, 1982–1999 e1988 (2021). 10.1016/j.stem.2021.07.011

35 Zhang, J. et al. Asparagine plays a critical role in regulating cellular adaptation to glutamine depletion. Mol Cell 56, 205–218 (2014). 10.1016/j.molcel.2014.08.018

36 Newsholme, E. A., Crabtree, B. & Ardawi, M. S. The role of high rates of glycolysis and glutamine utilization in rapidly dividing cells. Biosci Rep 5, 393–400 (1985). 10.1007/bf01116556

37 Jun, S. et al. The requirement for pyruvate dehydrogenase in leukemogenesis depends on cell lineage. Cell Metab 33, 1777–1792 e1778 (2021). 10.1016/j.cmet.2021.07.016

38 Lyu, J. et al. A glutamine metabolic switch supports erythropoiesis. Science 386, eadh9215 (2024). 10.1126/science.adh9215

39 Greenwood, D. L. et al. Acly Deficiency Enhances Myelopoiesis through Acetyl Coenzyme A and Metabolic-Epigenetic Cross-Talk. Immunohorizons 6, 837–850 (2022). 10.4049/immunohorizons.2200086

40 Umemoto, T. et al. ATP citrate lyase controls hematopoietic stem cell fate and supports bone marrow regeneration. EMBO J 41, e109463 (2022). 10.15252/embj.2021109463

41 Gothert, J. R. et al. In vivo fate-tracing studies using the Scl stem cell enhancer: embryonic hematopoietic stem cells significantly contribute to adult hematopoiesis. Blood 105, 2724–2732 (2005). 10.1182/blood-2004-08-3037

42 Morrison, S. J., Hemmati, H. D., Wandycz, A. M. & Weissman, I. L. The purification and characterization of fetal liver hematopoietic stem cells. Proc Natl Acad Sci U S A 92, 10302–10306 (1995). 10.1073/pnas.92.22.10302

43 Umemoto, T., Hashimoto, M., Matsumura, T., Nakamura-Ishizu, A. & Suda, T. Ca(2+)-mitochondria axis drives cell division in hematopoietic stem cells. J Exp Med 215, 2097–2113 (2018). 10.1084/jem.20180421

44 Sudo, T. et al. The endothelial antigen ESAM monitors hematopoietic stem cell status between quiescence and self-renewal. J Immunol 189, 200–210 (2012). 10.4049/jimmunol.1200056

45 Zhang, Z. et al. Hematopoietic stem cells activate a latent differentiation pathway to facilitate recovery after 5-fluorouracil-induced myeloablation. Dev Cell 61, 1044–1060 e1047 (2026). 10.1016/j.devcel.2026.02.003

46 Sarrazy, V. et al. Disruption of Glut1 in Hematopoietic Stem Cells Prevents Myelopoiesis and Enhanced Glucose Flux in Atheromatous Plaques of ApoE(-/-) Mice. Circ Res 118, 1062–1077 (2016). 10.1161/CIRCRESAHA.115.307599

47 Stoltzman, C. A. et al. Glucose sensing by MondoA:Mlx complexes: a role for hexokinases and direct regulation of thioredoxin-interacting protein expression. Proc Natl Acad Sci U S A 105, 6912–6917 (2008). 10.1073/pnas.0712199105

48 Wu, N. et al. AMPK-dependent degradation of TXNIP upon energy stress leads to enhanced glucose uptake via GLUT1. Mol Cell 49, 1167–1175 (2013). 10.1016/j.molcel.2013.01.035

49 Wilde, B. R., Ye, Z., Lim, T. Y. & Ayer, D. E. Cellular acidosis triggers human MondoA transcriptional activity by driving mitochondrial ATP production. Elife 8 (2019). 10.7554/eLife.40199

50 Pizzato, H. A. et al. Mitochondrial pyruvate metabolism and glutaminolysis toggle steady-state and emergency myelopoiesis. J Exp Med 220 (2023). 10.1084/jem.20221373

51 Fergestad, T., Bostwick, B. & Ganetzky, B. Metabolic disruption in Drosophila bang-sensitive seizure mutants. Genetics 173, 1357–1364 (2006). 10.1534/genetics.106.057463

52 Jaiswal, M. et al. Impaired Mitochondrial Energy Production Causes Light-Induced Photoreceptor Degeneration Independent of Oxidative Stress. PLoS Biol 13, e1002197 (2015). 10.1371/journal.pbio.1002197

53 Osawa, M., Hanada, K., Hamada, H. & Nakauchi, H. Long-term lymphohematopoietic reconstitution by a single CD34-low/negative hematopoietic stem cell. Science 273, 242–245 (1996). 10.1126/science.273.5272.242

54 Hinge, A. et al. Asymmetrically Segregated Mitochondria Provide Cellular Memory of Hematopoietic Stem Cell Replicative History and Drive HSC Attrition. Cell Stem Cell 26, 420–430 e426 (2020). 10.1016/j.stem.2020.01.016

55 Ardawi, M. S. & Newsholme, E. A. Glutamine metabolism in lymphocytes of the rat. Biochem J 212, 835–842 (1983). 10.1042/bj2120835

56 Altman, B. J., Stine, Z. E. & Dang, C. V. From Krebs to clinic: glutamine metabolism to cancer therapy. Nature Reviews Cancer 16, 619–634 (2016).

57 Sullivan, L. B. et al. Aspartate is an endogenous metabolic limitation for tumour growth. Nat Cell Biol 20, 782–788 (2018). 10.1038/s41556-018-0125-0

58 Potter, V. R., Le, P. G. & Klug, H. L. The assay of animal tissues for respiratory enzymes; oxalacetic acid oxidation and the coupled phosphorylations in isotonic homogenates. J Biol Chem 175, 619–634 (1948).

59 Potter, V. R. & Busch, H. Citric acid content of normal and tumor tissues in vivo following injection of fluoroacetate. Cancer Res 10, 353–356 (1950).

60 Kuhn, R., Schwenk, F., Aguet, M. & Rajewsky, K. Inducible gene targeting in mice. Science 269, 1427–1429 (1995). 10.1126/science.7660125

61 de Boer, J. et al. Transgenic mice with hematopoietic and lymphoid specific expression of Cre. Eur J Immunol 33, 314–325 (2003). 10.1002/immu.200310005

62 Mingote, S. et al. Genetic Pharmacotherapy as an Early CNS Drug Development Strategy: Testing Glutaminase Inhibition for Schizophrenia Treatment in Adult Mice. Front Syst Neurosci 9, 165 (2015). 10.3389/fnsys.2015.00165

63 Zhao, S. et al. ATP-Citrate Lyase Controls a Glucose-to-Acetate Metabolic Switch. Cell Rep 17, 1037–1052 (2016). 10.1016/j.celrep.2016.09.069

64 Chen, G. et al. Loss of vitamin C biosynthesis protects from the pathology of a parasitic infection. Proc Natl Acad Sci U S A 122, e2517730122 (2025). 10.1073/pnas.2517730122

65 Su, X., Lu, W. & Rabinowitz, J. D. Metabolite Spectral Accuracy on Orbitraps. Analytical Chemistry 89, 5940–5948 (2017). 10.1021/acs.analchem.7b00396

